# An Antibody-Based Molecular Switch for Continuous Biosensing

**DOI:** 10.1101/2023.03.07.531602

**Authors:** Ian A.P. Thompson, Jason Saunders, Liwei Zheng, Amani A. Hariri, Nicolò Maganzini, Alyssa P. Cartwright, Jing Pan, Michael Eisenstein, Hyongsok Tom Soh

## Abstract

We present a generalizable approach for designing biosensors that can continuously detect specific biomarkers in real time and without sample preparation. This is achieved by converting existing antibodies into target-responsive “antibody-switches” that enable continuous optical biosensing. To engineer these switches, antibodies are linked to a molecular competitor through a DNA scaffold, such that competitive target binding induces scaffold switching and fluorescent signaling of changing target concentrations. As a demonstration, we designed antibody-switches that achieve rapid, sample-preparation-free sensing of digoxigenin and cortisol in undiluted plasma. We showed that, by substituting the molecular competitor, we can further modulate the sensitivity of our cortisol switch to achieve detection at concentrations spanning 3.3 nM to 3.3 mM. Finally, we integrated this switch with a fiber-optic sensor to achieve hours-long continuous sensing of cortisol in buffer with <5-minute time resolution. We believe this modular sensor design can enable continuous biosensor development for many biomarkers.

## Introduction

Sensors capable of continuously measuring molecular analytes in a compact format offer tremendous potential in personalized health monitoring(*1*–*5*), early disease detection(*6, 7*), and inpatient care(*8*). A few such continuous biosensors have been developed to date, and now play a critical role in modern medicine. Continuous glucose monitors (CGMs) have revolutionized diabetes management by enabling long-term monitoring of a patient’s glycemic control and generating real-time alerts for rapid intervention(*9, 10*), and pulse oximetry has become an essential tool for directly measuring blood oxygenation during critical care(*11*). However, both of these technologies rely on analyte-specific sensing mechanisms; the former uses a glucose-specific enzyme that produces redox-active species,(*12*) whereas the latter relies on the intrinsic optical properties of oxygenated blood(*13*). As such, there is a considerable opportunity for more generalizable biosensor frameworks that can readily be adapted to recognize a broad spectrum of clinically important biomarkers. This will require the development of new continuous molecular detection strategies that can be rapidly and reliably adapted to many different biochemical targets.

Designing such a “universal” continuous biosensing mechanism presents a tremendous technological challenge. The sensing mechanism must achieve rapid, sensitive, and specific molecular recognition that generates a measurable signal output—ideally, using readily available affinity reagents as building blocks. To operate continuously at the point-of-care, sensing should also require minimal to no sample preparation or additional reagents to achieve detection. Switch-based biosensors have emerged as a promising solution that combines molecular recognition and signaling capabilities within a single receptor to enable continuous sensing without the need for additional reagents(*14*). To date, such biosensors have primarily employed sensor surfaces functionalized with DNA-based aptamer switches that change their conformation upon binding to a target, thereby enabling sensing through electrochemical or fluorescent reporters coupled to the DNA sequence. Aptamer switches have been applied to sensing a range of targets(*15*–*20*), but they remain difficult to generalize. This is partly because there is only a limited pool of high-performance aptamers available, such that each new aptamer-based sensor requires the selection of a new aptamer that binds the relevant analyte with excellent affinity and specificity(*21*). In many cases, these newly-selected aptamers must be further engineered to ensure that they undergo binding-induced conformational switching to generate a measurable signal(*22*). Both of these processes are laborious and unpredictable in terms of success rates, and thus the use of aptamer switch-based biosensors has remained limited to sensing a relatively small number of important biomarkers. As such, there remains an unmet need for robust and generalizable strategies for developing new molecular switches that can greatly expand the range of continuous biosensing applications.

To this end, we have developed a strategy for adapting existing antibodies into ‘antibody-switches’ that can achieve continuous molecular detection. Antibodies present an attractive alternative to aptamers as they are broadly available for a wide range of targets and offer a conserved, predictable structure that should make a generalizable engineering strategy feasible. In our antibody-switch design, an antibody is site-specifically augmented with a DNA scaffold to form a chimeric molecule that undergoes target-responsive switching upon antibody-antigen binding. This switching occurs through intramolecular competitive binding between the target antigen and a molecular “bait” linked to the end of the DNA scaffold. By introducing a Förster resonance energy transfer (FRET) fluorophore reporter pair within the antibody-switch, we can measure conformational changes in the switch through its fluorescence and thereby continuously measure changing target concentrations. We have validated our approach by engineering an antibody-switch that responds to the steroid digoxigenin. This antibody-switch achieved FRET-based measurement of changing concentrations across a wide dynamic range spanning ∼100 nM to 1 mM, with ∼5-minute temporal resolution and high sensitivity even in undiluted plasma. To illustrate the generalizability of this approach, we also engineered an antibody-switch for cortisol, which exhibits similarly robust performance. By tuning the identity of the bait molecule to a cortisol analogue with lower antibody affinity, we were able to further improve antibody-switch sensitivity by 100-fold. Finally, we incorporated our cortisol antibody-switch into an optical fiber biosensor design and demonstrate continuous cortisol sensing in buffer, with the ability to track fluctuating cortisol concentrations spanning the nanomolar to micromolar range with ∼5-minute temporal resolution for > 3 hours. We believe this modular design strategy should be broadly generalizable to virtually any existing antibody, thereby accelerating the proliferation of continuous biosensors for new targets.

## Results

### Design of an antibody-based molecular switch

In the antibody-switch design, an IgG antibody is engineered into a molecular switch that can achieve continuous biosensing of its cognate antigen with no sample preparation required. We use a DNA scaffold to link the antibody to a bait molecule—an analog of its antigen that competes with the binding of free antigen molecules present in the sample (**Figure 1a**). Switching occurs when the free antigen displaces the bound bait, or vice versa, leading to a conformational change in the DNA scaffold linking the bait and antibody. Target-dependent switching is quantified with high specificity through FRET-based fluorescent measurements derived from donor fluorophores on the surface of the antibody and an acceptor fluorophore incorporated within the DNA scaffold adjacent to the bait, such that the distance between donor-acceptor dye pairs increases when the antibody-switch opens in response to target binding (**Figure 1b**). This target-dependent switching is described by a three-state competitive switching equilibrium (**Supplementary Discussion 1**). In the absence of target, the linked bait molecule binds the antibody through an intramolecular reaction 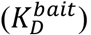. When the target is present, competition between this intramolecular reaction and antibody-target binding 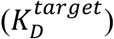 leads to switching of the construct from the closed (high FRET ratio) to the target bound (low FRET ratio) state.

**Figure 1.**
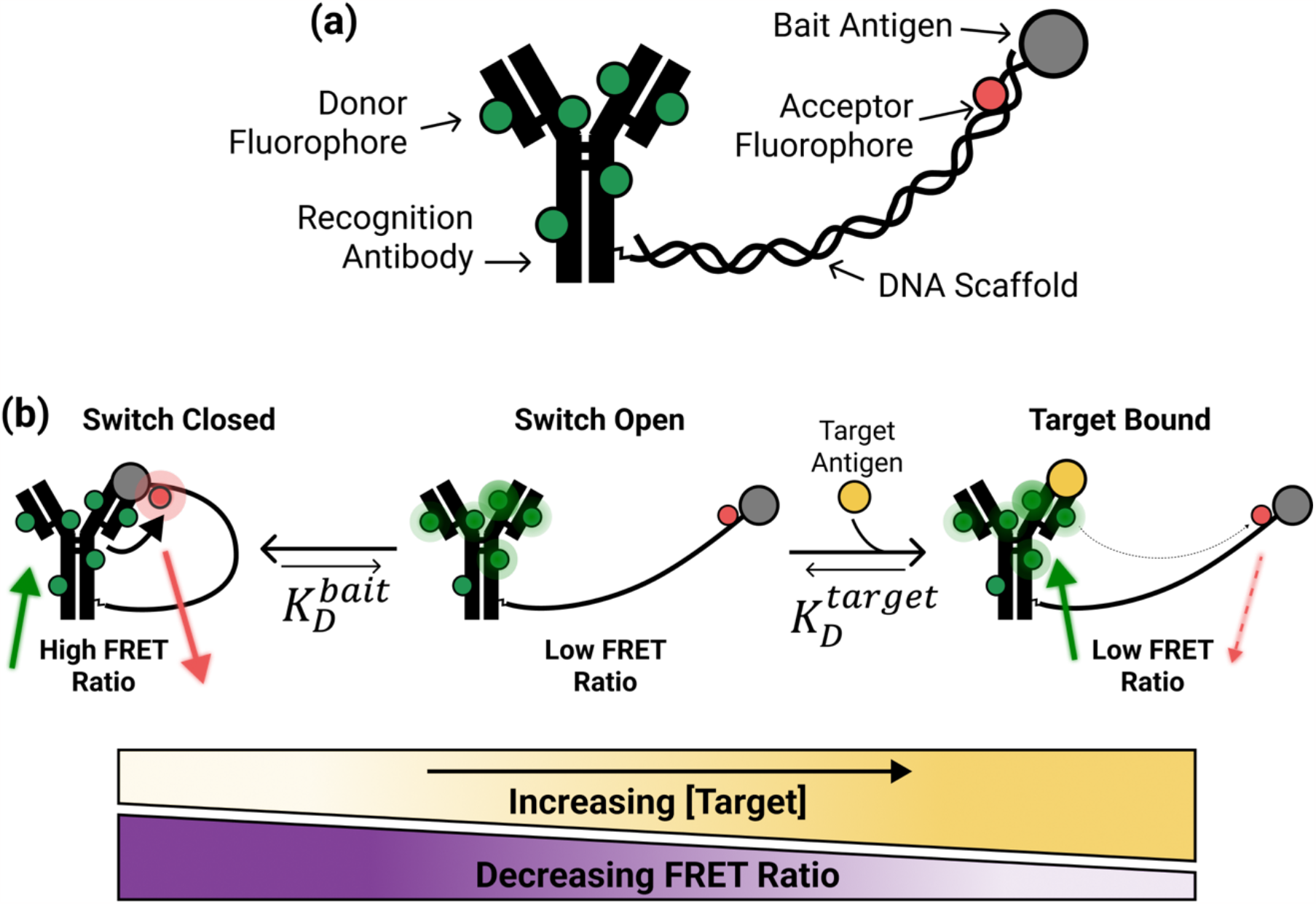
The antibody-switch construct. **(a)** We adapt an antibody into a molecular switch by tethering it to a bait molecule through a DNA scaffold. Fluorescent quantification of changing target analyte concentrations is achieved by incorporating FRET donor and acceptor fluorophores on the antibody surface and within the DNA scaffold, respectively. **(b)** The molecular switch operates through a three-state competitive equilibrium, where increased target concentrations shift the equilibrium towards the target-bound (right) and away from the closed state (left). This equilibrium depends on the target affinity of the antibody 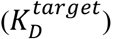, but target sensitivity can also be tuned by varying the strength of intramolecular bait binding 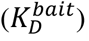. The switch exhibits a decreased FRET ratio upon target binding as the distance between the donor and acceptor fluorophores is increased in the open and bound states compared to the closed state.

Although the affinity of the chosen antibody is fixed, we can tune the strength of the intramolecular reaction to control the overall sensitivity of our switch(*23*). If the bait molecule is identical to the target antigen, intramolecular bait binding will be strong (low 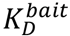), shifting the equilibrium towards the closed state and resulting in low target sensitivity. Low sensitivity here is defined by a high EG_50_, which describes the free target concentration at which a half-maximal FRET switching response occurs. In contrast, a mismatched bait molecule with a higher 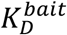 leads to weaker competition and a more target-sensitive switch (low EC_50_). However, if antibody-bait binding is too weak, intramolecular binding will not occur, leading to high background from a large proportion of switches in the open but non-target-bound state and thus a poor signal-to-noise ratio. It is also critical to ensure that the bait and free target binding are fully competitive, such that displacement of the bait is guaranteed upon target binding. This means that careful selection of the bait antigen is needed to balance strong FRET signal and sensitivity to low target concentrations. Critically, the modular antibody-switch design can be applied to different small molecule targets by substituting the antibody-bait pair while maintaining a consistent DNA scaffold geometry and switch assembly process, as described in detail below.

### Synthesis and validation of a digoxigenin-sensing antibody-switch

As an initial proof of concept, we developed an antibody-switch that recognizes digoxigenin (DIG), a plant-derived steroid molecule. We first synthesized our DIG-DNA conjugate by covalently coupling DIG to the 5’ end of a 20-nt single-stranded bait-DNA oligonucleotide modified with a Cy5 acceptor dye molecule 10 nt from its 5’ end (**Figure 2a**; see **Supplementary Table 1** for all DNA sequences used in this work). The selection of DIG as both the target and the bait molecule should yield a switch exhibiting strong binding between the DIG-DNA and free target (low 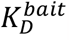) at the expense of lower switch sensitivity (high EC_50_).

**Figure 2.**
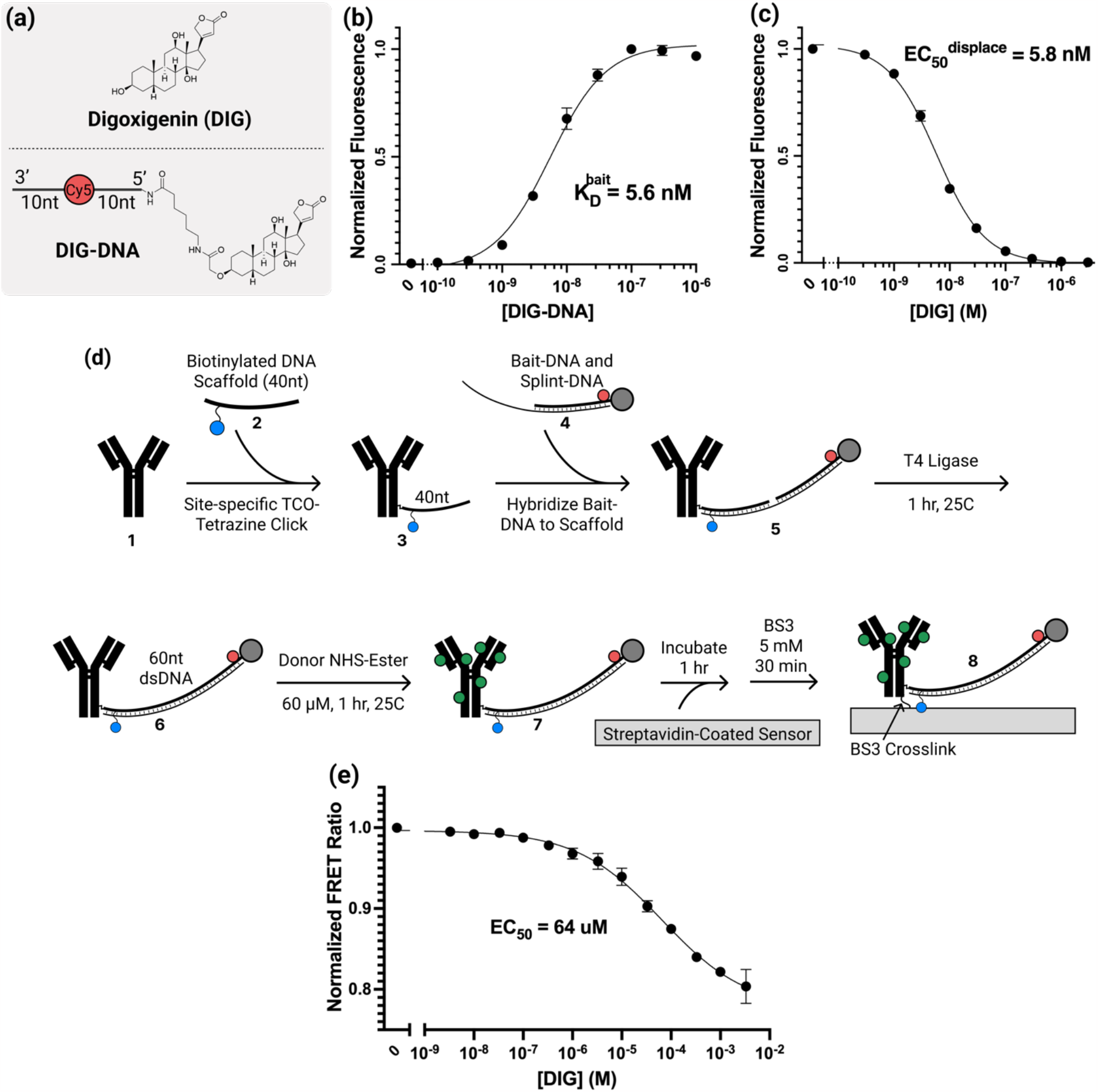
Engineering a DIG-responsive antibody-switch. **(a)** Our DIG-DNA construct incorporates a DIG bait moiety and a Cy5 fluorescent reporter into a 20-nt DNA scaffold. **(b)** We used flow cytometry to characterize the 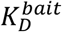 of DIG-DNA binding to a polyclonal anti-DIG antibody (pAb). **(c)** Competitive binding assays quantifying the disruption of DIG-DNA/pAb antibody pair binding by free DIG in solution suggest that the molecular competition from DIG-DNA can be used to quantify target binding. We used 10 nM DIG-DNA for all measurements in this plot. **(d)** The multi-step antibody-switch synthesis process leverages site-specific antibody-DNA conjugation followed by splinted ligation to assemble a complete switch with controlled stoichiometry and a site-specific biotin handle for immobilization onto biosensor surfaces. **(e)** When assembled antibody-switches are assembled on magnetic beads and measured using flow cytometry, the antibody-switch FRET signaling response spans a broad dynamic range of DIG concentrations. All datapoints and error bars represent the mean and standard deviation of three replicates.

To quantify DIG-DNA/antibody affinity, we incubated magnetic beads coated with candidate antibodies with varying concentrations of free (not antibody conjugated) DIG-DNA, washed, and then analyzed the beads via flow cytometry to quantify the bound fraction of DIG-DNA. Because the steric impact of DNA conjugation on DIG-antibody binding is hard to predict, we tested two different antibodies—one polyclonal and one monoclonal—to ensure that this conjugation did not disrupt the competitive binding needed for proper antibody-switch function. We found that the polyclonal antibody (pAb) exhibited superior properties for constructing an antibody-switch, binding to DIG-DNA with a 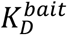 of 5.6 nM (95% CI = 4.8–6.4 nM), indicating that the attachment of DNA did not dramatically disrupt the antibody-binding interface (**Figure 2b**). We next incubated pAb-coated beads with a fixed concentration of DIG-DNA and varying concentrations of free target and measured the bound fraction of DIG-DNA that was not displaced by free DIG. Notably, free DIG robustly outcompeted DIG-DNA; with 10 nM DIG-DNA, a DIG concentration of 5.8 nM (95% CI = 5.4–6.2 nM) was sufficient to displace half of the bound DIG-DNA, indicating that free DIG exhibited slightly stronger antibody binding than DIG-DNA (**Figure 2c**). In contrast, when we performed the same analysis with the monoclonal antibody, DIG-DNA exhibited weaker affinity, with a 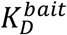 of 91 nM (95% CI = 56–148 nM) and poor competitive displacement by the free target: > 5 μM free DIG was required to achieve 50% DIG-DNA displacement with 10 nM DIG-DNA (**Supplementary Figure 1**). This poorer performance was likely due to steric interference of the DNA within the DIG-DNA conjugate with the monoclonal antibody’s capacity to recognize DIG in solution, and we therefore selected pAb for subsequent experiments.

We subsequently synthesized the complete antibody-switch construct (**Figure 2d**). Starting from unmodified pAb **1**, we site-specifically modified the glycans on the antibody Fc domain, and then conjugated these sites with a 40-nt scaffold-DNA **2** that bears a biotin moiety 6 nt from the antibody conjugation site for purposes of surface immobilization. Such site-specific conjugation ensures consistency in terms of the antibody-switch geometry, which in turn dictates intramolecular competition and thus switching function. Any unconjugated scaffold-DNA was then removed by size-exclusion purification of the antibody-DNA conjugate. Using SDS-PAGE, we determined that the resulting antibody-DNA conjugates **3** exhibit ∼1 DNA molecule per Fc domain, as intended (**Supplementary Figure 2**). Next, we coupled the DIG-DNA complex to the antibody-DNA conjugate using enzymatic ligation. We first hybridized DIG-DNA with a partially complementary scaffold-complement sequence, where the overhang is complementary to the pAb-coupled scaffold-DNA sequence **4**. This assembly then forms the splinted 60-bp DNA scaffold for the antibody-switch, with a nick between the DIG-DNA strand and the scaffold-DNA strand **5**. Incubation with T4 ligase subsequently closes this nick **6** (**Supplementary Figure 3**). Donor fluorophores (Alexa Fluor 546) were then incorporated by modifying the antibody’s exposed lysine residues **7**. The complete switch can then be site-specifically functionalized onto a streptavidin-bearing biosensor surface via biotin-streptavidin interaction, which is reinforced by adding a BS3 amine-to-amine crosslinker to achieve covalent surface attachment **8**. Critically, this synthesis process ensures, through purification of the antibody-DNA conjugate and the high efficiency of ligation, that only fully assembled antibody-switch constructs contain the biotin moiety required for surface immobilization. This modular assembly process also allows us to substitute different bait molecules, DNA scaffolds, or antibody candidates to modify switch function.

We confirmed that our antibody-switch undergoes target-responsive FRET switching by immobilizing the switches onto streptavidin-coated magnetic beads and then incubating the beads for 1 hour with various concentrations of DIG in a buffer that was selected to match ionic concentrations in blood and interstitial fluid (ISF)(1x PBS with 1 mM MgCl_2_ and 4 mM KCl). We measured the fluorescent response via flow cytometry immediately afterward. As expected, the FRET ratio decreased as we introduced increasing DIG concentrations **(Figure 2e, Supplementary Figure 4a)**. We fitted the FRET response to a Hill binding model and determined that the sensor responds with a Hill coefficient (n) of 0.51 (95% CI = 0.44–0.59) and an EC_50_ of 64 μM (95% CI = 35–93 μM). The switch responded to changes in DIG concentration spanning a greater than 10,000-fold dynamic range, with an upper bound in the low-millimolar range and a limit of detection (LOD) of 68 nM.

### Specific, rapid, and reversible antibody-switch biosensing

We next assessed our antibody-switch’s suitability for continuous biosensing in complex biofluids, where nonspecific binding by interferents poses a major challenge. We assembled antibody-switch-coated beads as described above and then tested their specificity by challenging them with DIG doped into the protein-rich environment of undiluted chicken plasma and then measuring their FRET response via flow cytometry. Before measurement, plasma samples were filtered with a 0.45 μM syringe filter to remove large particulates that interfere with flow cytometry while preserving the complete protein, small molecule, and fluid content of the plasma. Remarkably, the switch response in undiluted plasma was virtually identical to that in buffer, with an identical LOD of 68 nM and a largely unchanged dynamic range, where EC_50_ = 87 μM (95% CI = 15–160 μM) and n = 0.43 (**Figure 3a, Supplementary Figure 4b**).

**Figure 3.**
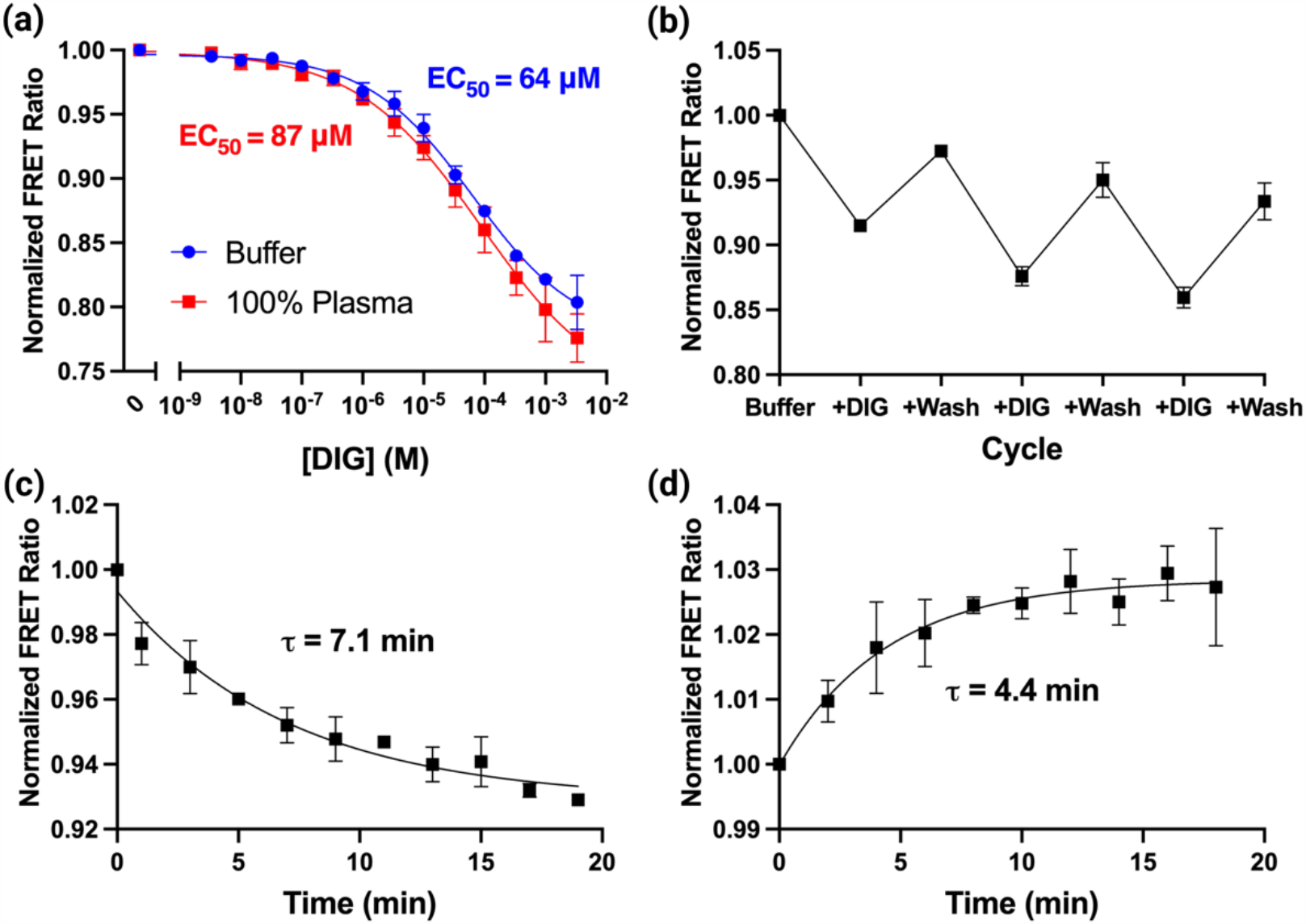
Assessing suitability of the DIG antibody-switch for continuous sensing in complex media. **(a)** Our antibody-switch achieved essentially identical DIG-responsive FRET sensing performance in both buffer and undiluted chicken plasma when immobilized on magnetic beads and measured using flow cytometry. **(b)** Reversible binding and signaling over multiple 30-minute cycles of target addition and removal indicates that our antibody-switch is suitable for repeated or continuous measurements. We observed a rapid kinetic response from our antibody-switch, with an **(c)** association time constant of 7.1 min (95% CI = 3.4–8.0 min) and **(d)** a dissociation time constant of 4.4 min (95% CI = 2.8–7.5 min). All datapoints and error bars represent the mean and standard deviation of three replicates.

The ability to sense changes in target concentration rapidly and reversibly is also crucial for continuous sensors, and we hypothesized that our linked, intramolecular design would enable rapid switching between open and closed states over many cycles of changing target concentrations. We repeatedly exposed our bead-immobilized antibody-switches to high concentrations of DIG (1 mM) in buffer for 30 min, followed by 30-min incubations in target-free buffer. We observed reversible and repeatable FRET switching over the course of a three-hour experiment involving three cycles of target addition and wash-out (**Figure 3b**). We observed a small degree of FRET ratio decrease over each cycle, and we hypothesized that this may be due to residual DIG that remains after wash cycles.

We also measured the antibody-switch’s FRET response kinetics upon DIG addition and washing with buffer. We observed rapid changes in FRET ratio in terms of both target binding (τ_on_ = 7.1 min, **Figure 3c**) and dissociation (τ_off_ = 4.4 min, **Figure 3d**). This indicates suitable performance for collecting continuous measurements with a temporal resolution of ∼5 minutes— and potentially faster, if sensor response is characterized with a pre-equilibrium approach(*24*). The roughly symmetric kinetic response to increasing or decreasing target concentrations is consistent with our competitive binding model, where both on- and off-switching events require the slow dissociation of either the free target or the bait (**Supplementary Discussion 1**).

### Engineering a cortisol-responsive antibody-switch with enhanced sensitivity using a mismatched bait design

As described above, our antibody-switch design can be readily tailored for new target biomolecules by simply substituting the antibody-bait pair, and we demonstrated this capability by engineering an antibody-switch for cortisol. Cortisol is an important stress hormone that affects nearly every organ system in the body, playing a critical role in stress, metabolism, and inflammation(*25, 26*). We hypothesized that by employing a cortisol analog as a bait molecule, we could tune the intramolecular competition within our switch—and thus its affinity for free cortisol—to achieve sensitive detection over clinically-relevant cortisol concentrations, which span approximately 1 nM to 1 μM in blood and ISF(*27*). We chose a cortisol-binding monoclonal antibody, and then screened antibody binding and competitive displacement for bait-DNA conjugates incorporating either cortisol or corticosterone (a cortisol analog with a missing hydroxyl group near position 21, where antibody recognition occurs) as the bait molecule (**Figure 4a**). The bait-DNA incorporating cortisol (Cort-DNA) exhibited a low 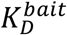 of 1.8 nM (95% CI = 1.6–2.0 nM; **Figure 4b, blue**) and competed strongly with free cortisol, with 41 nM cortisol (95% CI = 30–56 nM) required to achieve 50% displacement of bound Cort-DNA in the presence of 10 nM bait-DNA (**Figure 4c, blue**). In contrast, the corticosterone bait-DNA (CSO-DNA) exhibited a higher 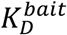 of 4.7 nM (95% CI = 3.7–6.0 nM; **Figure 4b, green**) and approximately 7-fold weaker competition with free cortisol, with only 5.8 nM (95% CI = 5.3–6.3 nM) of free cortisol required to achieve equivalent displacement of 10 nM CSO-DNA (**Figure 4c, green**).

**Figure 4.**
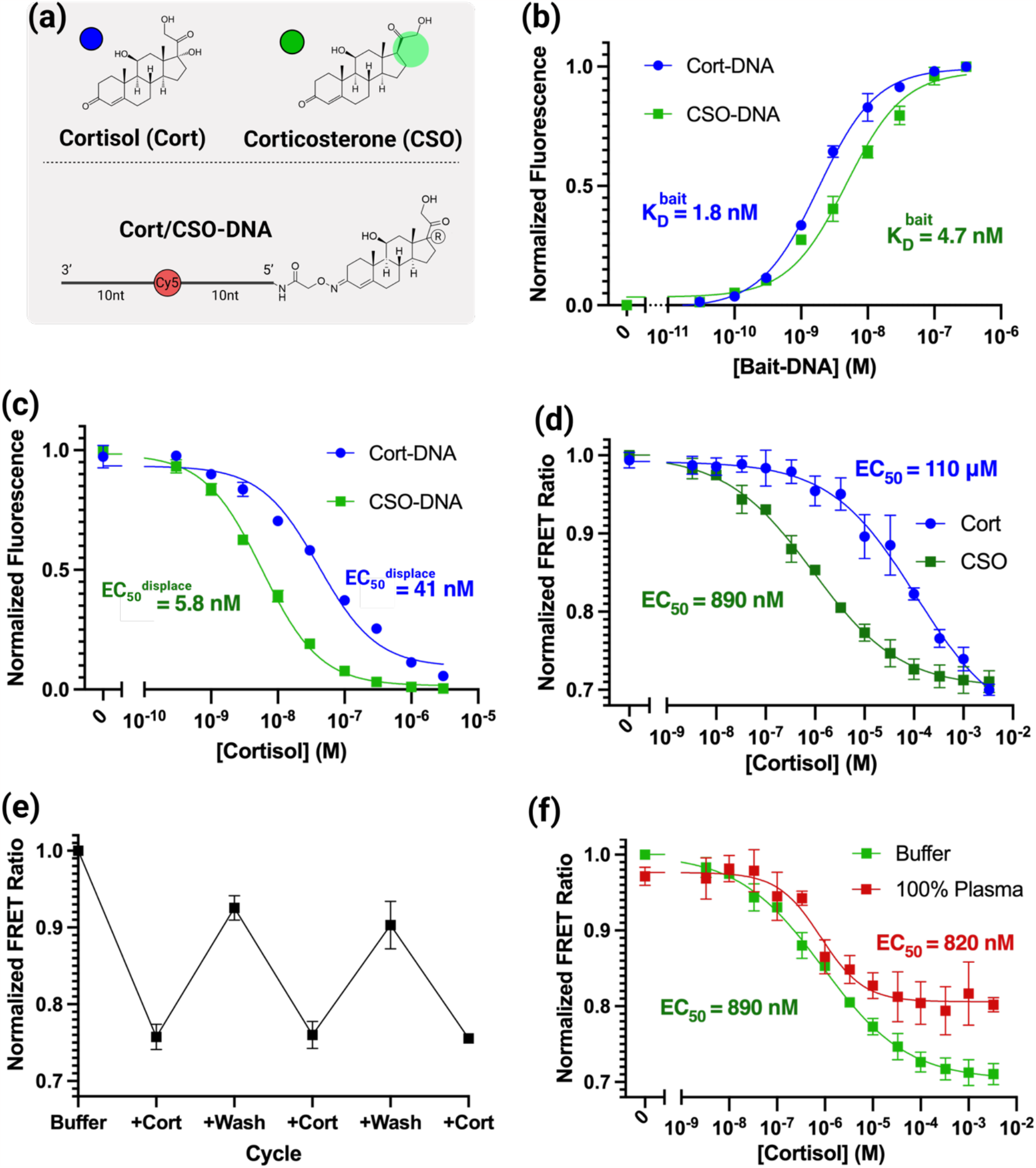
Cortisol antibody-switch engineering. **(a)** Our cortisol antibody-switch incorporated either cortisol (Cort-DNA) or corticosterone (CSO-DNA) as the bait molecule at the end of the 20-nt DNA scaffold. **(b)** Monoclonal anti-cortisol binding to bait-DNA conjugates. When CSO-DNA is used in place of Cort-DNA, we measured a higher 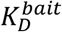 in bead-based antibody binding assays, indicating that corticosterone may act as a weaker competitor molecule. **(c)** Competitive binding assays of bait-DNA conjugates against free cortisol further demonstrate that CSO-DNA acts as a weaker competitor than Cort-DNA. All experiments were conducted with 10 nM bait-DNA. **(d)** Cortisol-dependent FRET responses of antibody-switches incorporating Cort-DNA or CSO-DNA in buffer. Both switches respond to a wide dynamic range of cortisol concentrations, but the CSO-DNA construct achieves higher sensitivity due to weaker intramolecular competition. **(e)** The CSO-DNA antibody-switch exhibits reversible binding over multiple cycles of target addition and washing. **(f)** This switch also maintains robust cortisol sensing in undiluted chicken plasma. All datapoints and error bars represent the mean and standard deviation of three replicates.

We followed the modular synthesis process described above to construct antibody-switches that combine XM210 with either Cort-DNA or CSO-DNA (**Supplementary Figure 2, 3b**), and then immobilized these onto streptavidin beads. Both switches exhibited decreasing FRET ratios at increasing target concentrations, indicating successful assembly of cortisol-responsive molecular switches (**Figure 4d; Supplementary Figure 5a, b**). The Cort-DNA switch achieved modest cortisol sensitivity, spanning from a LOD of 340 nM up to a maximum tested concentration of 3.3 mM. In comparison, the mismatched corticosterone bait enhanced antibody-switch sensitivity by ∼100-fold, with a LOD of 3.3 nM and an upper limit of quantification of 95 μM. This means that the CSO-DNA switch is likely to be better suited for monitoring clinically relevant basal cortisol concentrations. These results highlight how the selection of alternative analogs as bait can enable further optimization of sensor performance for a given analyte.

Like the DIG-responsive antibody-switch, the CSO-DNA switch exhibited robust reversibility and specificity in complex biofluids. When exposed to multiple 30-min cycles of 1 mM cortisol and wash-out with target-free buffer, the bead-immobilized CSO-DNA antibody-switches exhibited stable and reversible on-and-off switching (**Figure 4e**). We also tested our antibody-switch in chicken plasma, which shares a similar level of protein content to human plasma. However, since chickens primarily produce corticosterone rather than cortisol, we can exclude the possibility that changes in switching behavior arise from endogenous rather than spiked cortisol(*28*). The cortisol-responsive switch maintained its switching behavior and ability to sense spiked in cortisol with good sensitivity in undiluted plasma, with a LOD of 140 nM (**Figure 4f; Supplementary Figure 5c**). This is lower than the sensitivity we observed in buffer, most likely due to high levels of cortisol binding to plasma proteins such as albumin and transcortin, which may attenuate the free (non-protein-bound) levels to only 5–25% of the spiked-in cortisol concentration(*27, 29*). However, since the free fraction of cortisol is the only bioactive fraction, this may not impede our switch’s applicability for diagnostic purposes(*30*). These results thus demonstrate the generalizability of the antibody-switch design as a tool for achieving continuous biosensing in complex matrices.

### Continuous cortisol biosensing with antibody-switches

Finally, we assessed the continuous sensing performance of our antibody-switch when incorporated with an optical fiber-based biosensor described previously by our group(*31*). This system comprises a tapered optical fiber sensor functionalized with a monolayer of antibody-switches assembled onto the fiber tip surface via streptavidin-biotin immobilization (**Figure 5a, inset**). To assemble this sensor, tapered optical fibers were coated in a polyethylene glycol (PEG) passivation layer bearing a limited number of biotin end-groups (1% of the monolayer). These biotin groups were then coupled to the biotinylated antibody-switches via streptavidin molecules. The tapered profile of the fiber optic tip leads to the formation of an evanescent field of light excitation that is confined close to the sensor surface, only penetrating a few hundred nanometers into solution. This evanescent sensing strategy enables selective excitation of donor dyes within the antibody-switch without eliciting meaningful background from unbound molecules in the sample. It also enables efficient collection of emitted light from both donor and acceptor dyes for FRET ratio measurements. This optical fiber sensor is placed within a ∼50 μL sample chamber that allows us to flow samples across the sensor surface (**Figure 5a**). An optical reader couples laser excitation light into the optical fiber sensor, and then collects and measures the emitted light from antibody-switches immobilized onto the sensor surface. This enables continuous measurement of the sensor FRET response as target concentrations change.

**Figure 5.**
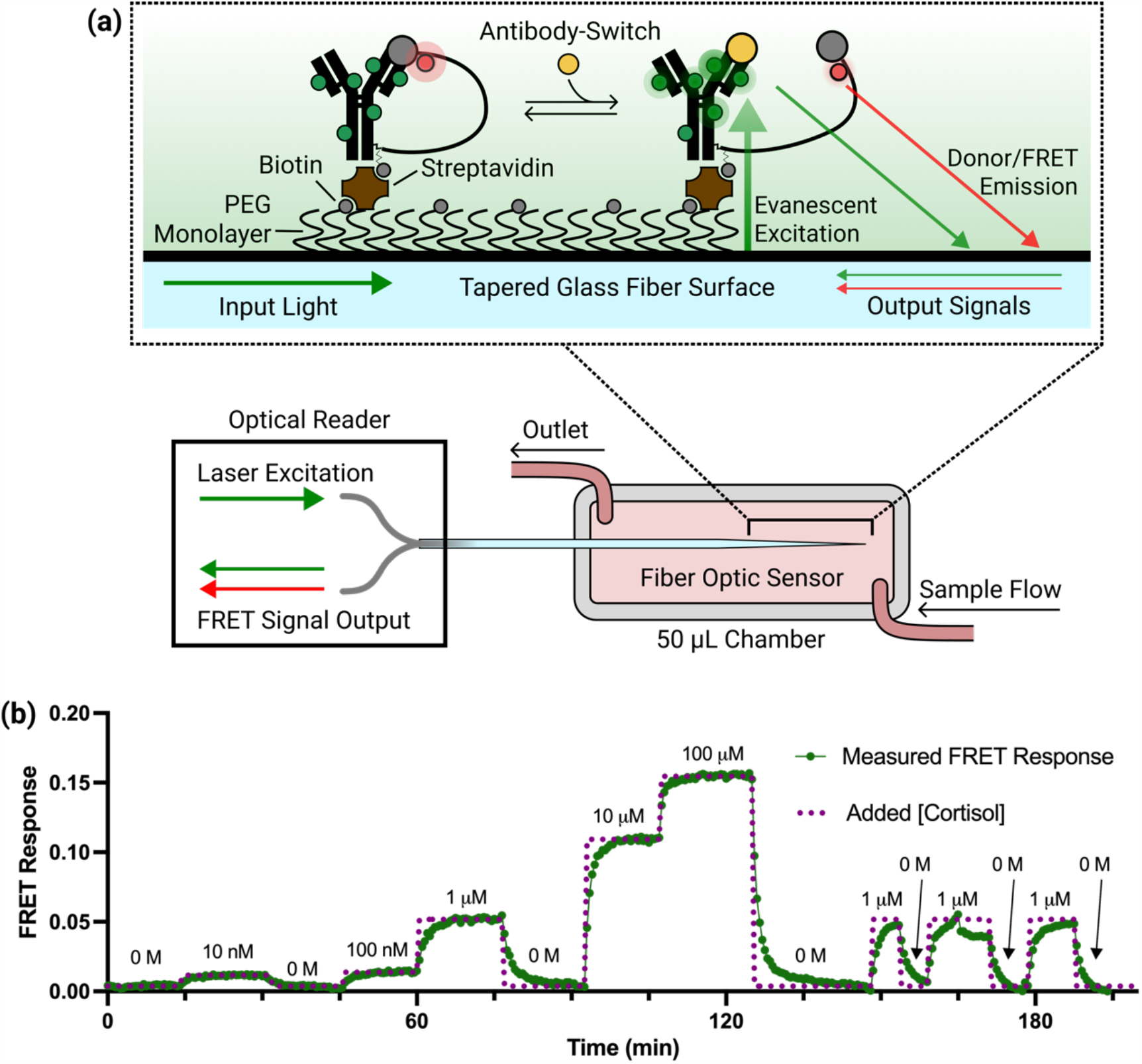
Continuous cortisol sensing with antibody-switches coupled to a fiber optic sensor. **(a)** Diagram of the setup for our tapered fiber optic sensor. The fiber tip is functionalized with antibody-switches via biotin-streptavidin immobilization onto a PEG monolayer. Laser light travels to the fiber tip and excites antibody-switch donor fluorophores within the evanescent field generated along the tapered portion of the fiber. The resulting donor and FRET emission is then coupled back to measurement instrumentation through the fiber. The measurements themselves are performed within a 50 μL flow chamber. **(b)** Continuous FRET measurement of changing cortisol concentrations in buffer over the course of a 3+ hour experiment. Buffer spiked with various concentrations of cortisol (purple dotted curve) was injected into the chamber at 5-to-15-minute intervals, and the antibody-switch FRET response was measured continuously. The presented FRET response data is corrected for the effects of photobleaching and plotted as the absolute value of the FRET ratio change from baseline so that increased concentrations lead to increased rather than decreased signals.

For continuous sensing experiments, we substituted the Cy5 acceptor dye with ATTO643, which exhibits better photobleaching resistance. To allow orthogonal modification of the bait-DNA with both our bait molecule and this enhanced dye, we used copper-free click chemistry to attach ATTO643 to an alkyne modification situated four bases away from the bait molecule (see ‘bait-DNA-Alkyne4’ in **Supplementary Table 1**). We moved the dye molecule closer to the bait in this design in an effort to increase FRET in the closed state and thus increase the FRET ratio change upon switching. Substituting the acceptor dye did not produce any substantial changes in the switching affinity of the Cort-DNA-Alkyne4 or CSO-DNA-Alkyne4 antibody-switches and moving the dye to a position 4 nt from the bait increased the binding-induced FRET response in the CSO-DNA-Alkyne4 switch (**Supplementary Figure 6a, b**). In addition, ATTO643 exhibited ∼20-fold slower photobleaching when exposed to continuous laser illumination compared to Cy5 (**Supplementary Figure 6c**). To further increase the long-term fluorescent stability of our sensor, we used intermittent laser excitation, acquiring measurements every 30 seconds. Because even slow photobleaching effects accumulate over time, data were corrected by first acquiring a photobleaching calibration measurement over 15 minutes in the absence of target, and then subtracting this calibrated photobleaching slope over the duration of measurement (**Supplementary Figure 7**).

We demonstrated the performance of our CSO-DNA-Alkyne4 antibody-switch-coupled fiber optic sensor over a series of stepwise cortisol concentration changes. We injected buffer spiked with varying cortisol concentrations into the measurement chamber to vary the free target concentration seen by the sensor at regular intervals, then allowed the sensor to equilibrate in the solution without flow while continuously measuring the FRET response. **Figure 5b** shows the FRET response over > 3 hours of continuous measurement, plotted as the inverse of the FRET ratio change such that increased concentrations are represented by increasing rather than decreasing signals. As in our bead-based assays, the switch was sensitive to sharp increases and decreases in cortisol concentration across a ∼10,000-fold dynamic range, detecting spikes as low as 10 nM and as high as 100 μM. Over the course of this measurement, the sensor responded rapidly to rising and falling cortisol concentrations, achieving equilibration in < 5 min at all measured concentrations (**Supplementary Figure 8**). Critically, the sensor maintained a stable baseline over the course of many cycles of increasing and decreasing target concentration.

## Discussion

The antibody-switch design provides a way to engineer existing antibodies into target-responsive molecular switches capable of real-time continuous biosensing. Our constructs link an antibody to a bait molecule through a DNA scaffold, enabling competitive binding with the free target that gives rise to conformational switching events that can be detected via FRET. These switches exhibit desirable properties for biosensing, including a broad (∼10,000-fold) dynamic range, rapid kinetics, reversibility, and the ability to sense targets directly in undiluted biofluids. The antibody-switch design is also modular and generalizable, such that it can be tailored to a broad range of targets simply by exchanging the antibody and bait molecule, as we have demonstrated in experiments using antibody-switches for DIG and cortisol. This modularity also allowed us to tune the sensitivity of our cortisol-responsive antibody-switch by replacing the cortisol bait with a mismatched corticosterone bait, which enhanced the switch’s target affinity ∼100-fold. By integrating our switch into a miniaturized fiber optic sensor probe, we were able to achieve continuous cortisol sensing across a concentration range of 10 nM to 100 μM with temporal resolution of < 5 minutes and stability over > 3 hours in buffer. These preliminary results suggest that antibody-switches may offer a general and robust method for developing switch-based continuous biosensors.

Various groups have achieved reagent-free continuous biosensing of large protein targets using protein-based molecular switches. However, these sensing strategies are reliant upon substantial changes in receptor mass upon binding(*32*), folding responses that are intrinsic to specific protein receptors(*33*), or two-site binding to a large protein target(*34*), and thus are not amenable to sensing small molecules. Our antibody-switch complements these technologies by enabling continuous detection of important small molecule biomarkers like cortisol. This design also offers improved temporal resolution relative to the abovementioned approaches, with sensor responses to changing analyte concentrations occurring within minutes, as compared to the ∼1-hour timescales achieved by other protein-based sensing strategies(*32*). We also anticipate that future iterations of the design could accommodate equally rapid continuous biosensing of larger protein targets by adapting the DNA scaffold design to accommodate larger protein fragments as bait molecules.

Our design provides a general strategy for transforming virtually any antibody into a monolithic sensor construct that achieves target recognition, molecular switching, and fluorescent signaling with only minimal screening effort required and without any engineering of the antibody protein itself. This approach thus eliminates the need for unique proteins with intrinsic switching, high-throughput combinatorial selection assays, or case-by-case redesign of the switch structure. As such, we believe that this antibody-switch approach has the potential to accelerate development of a diverse range of continuous biosensors for physiological monitoring and improved patient care.

## Materials and Methods

### Reagents & Materials

All oligonucleotide sequences (**Supplementary Table 1**) were purchased HPLC-purified from Integrated DNA Technologies with various internal and terminal modifications as specified in their sequences. Cortisol monoclonal antibody (XM210) was obtained from Abcam. Digoxigenin polyclonal antibody (Sheep, SKU: 11333089001) was obtained from Sigma Aldrich and digoxigenin monoclonal antibody (Clone # 611621) was obtained from R&D Systems. T4 Ligase and low molecular-weight DNA ladder were obtained from New England Biolabs (NEB). Methyltetrazine DBCO, TCO-NHS Ester (Axial), and BTTES were purchased from Click Chemistry Tools. ATTO643 azide was purchased from ATTO-Tech GmbH. VECTABOND reagent was purchased from Vector Laboratories. All DNA synthesis reagents, including 5’-amino-modifier C6, alkyne-modifier serinol phosphoramidites, and Glen-Pak DNA purification cartridges were purchased from Glen Research. Biotin-PEG-SVA (MW 5,000) and mPEG-succinimidyl valerate (MW 5,000) were purchased from Laysan Bio. Amicon Ultra-0.5 size exclusion columns, digoxigenin, digoxigenin NHS-ester (DIG-NHS), hydrocortisone, hydrocortisone 3-(O-carboxymethyl)oxime, O-(carboxymethyl)hydroxylamine hemihydrochloride, dimethylformamide (DMF), dimethylsulfoxide (DMSO), sodium ascorbate, and copper (II) sulfate were purchased from Sigma Aldrich. Cortisol-3CMO (O-(Carboxymethyl)hydroxylamine hemihydrochloride)), 1-ethyl-3-(3-dimethylaminopropyl)carbodiimide hydrochloride (EDC), N-hydroxysulfosuccinimide (NHS), Alexa Fluor 546 NHS ester (succinimidyl ester), Dynabeads protein G, Dynabeads T1, SiteClick antibody azido modification kits, bis(sulfosuccinimidyl)suberate (BS3), Tris-HCL (1 M, pH 7.5), magnesium chloride, 10x PBS (pH 7.4), ATP, NuPAGE sample reducing agent, NuPAGE LDS Sample Buffer (4x), NuPAGE 4 to 12% Bis-Tris mini protein gels, 20x NuPAGE MES SDS running buffer, SeeBlue pre-stained protein standard, SimplyBlue SafeStain, UltraPure TBE buffer (10X), Novex TBE-Urea Gels (10%), GelStar nucleic acid gel stain, and Tween 20 were purchased from Thermo Fisher Scientific.

### Instrumentation

Standard automated oligonucleotide solid-phase synthesis was performed on an Expedite 8909 synthesizer from Biolytic Lab Performance. DNA was quantified by UV absorbance at 260 nm with the NanoDrop 2000 spectrophotometer from Thermo Fisher Scientific. Gel electrophoresis was carried out on a 20 × 20 cm vertical Hoefer 600 electrophoresis unit. Gel images were captured using a ChemiDoc MP System from Bio-Rad Laboratories. Thermal annealing of all DNA structures was conducted using an Eppendorf Mastercycler 96-well thermocycler. Flow cytometry was performed using an SH800 cell sorter from Sony Biotechnology. Mass spectrometry (MS) was performed using a Microflex Matrix-assisted laser desorption/ionization time-of-flight (MALDI-TOF) instrument from Bruker. Shaking was done using a ThermoMixer F1.5 from Eppendorf.

### Digoxigenin bait-DNA Synthesis

Digoxigenin bait-DNA conjugates were prepared by labeling the amine in the bait-DNA sequence with DIG-NHS. DIG-NHS was dissolved in 15 mM DMF and then added to the bait-DNA in 100 mM sodium bicarbonate (pH 9) to achieve final concentrations of 82 μM DNA and 1.2 mM DIG-NHS in a 122 μL reaction volume. This mixture was incubated with rotation overnight at room temperature, then purified via ethanol precipitation and resuspended in DI water to ∼100 μM before use. MS was used to validate DNA modification.

### Testing binding and competition for bait-DNA candidates

Pairs of target-binding antibodies and bait-DNA conjugates were screened to assess their affinity and the ability of free target to compete with their binding. All assays were performed in physiological buffer (1x PBS (pH 7.4) with 1 mM MgCl_2,_ 4 mM KCl, and 0.05% Tween-20). The antibodies were first immobilized onto Dynabeads protein G by diluting 6 μL of stock Dynabeads (30 mg/mL) into 600 μL of 230 nM antibody in buffer. Beads were incubated for 1 hour and then washed and resuspended in 6 mL of buffer. Samples for each bait-DNA/target concentration were prepared by mixing 90 μL of antibody-coated beads with 10 μL of bait-DNA/target solution to achieve the final desired concentrations. For the bait-DNA binding assays, no target was included, and the concentration of bait-DNA varied between 0–3 μM. For competition assays, bait-DNA concentrations were held constant at 10 nM and free digoxigenin concentrations (stock prepared at 100 mM in DMSO) were varied between 0–100 μM. Antibody-coupled beads were incubated in these solutions for 1 hour, then resuspended in 200 μL of buffer immediately before measuring on a cell sorter to quantify Cy5 fluorescence.

### Cortisol and corticosterone bait-DNA synthesis

The Cort-DNA conjugate was prepared by coupling cortisol-3CMO to the terminal amine on the bait-DNA sequence using EDC/NHS chemistry. First, we combined 50 μL of 100 mM steroid-3CMO conjugate in 100% DMSO, 50 μL of 100 mM sulfo-NHS in 50% DMSO, and 5 μL of 100 mM EDC in 100% DMSO, and then shook this mixture at 1,500 rpm for 30 minutes at room temperature using a thermomixer. We then added 100 μL of 100–500 μM DNA and 40 μL of 1M sodium bicarbonate (pH 9) and incubated this mixture for 2 hours at room temperature. The DNA was buffer exchanged into DI water using a 3 kDa MWCO Amicon Ultra-0.5 centrifugal filter, and then purified using HPLC and validated for correct bait conjugation via MS.

CSO-DNA was synthesized by adapting a previously-described protocol(*35*). A carboxyl-group was conjugated onto corticosterone by combining 1.480 mL of 75 mM carboxymethoxylamine hemihydrochloride in methanol, 1.184 mL of 125 mM corticosterone in methanol, and 23.7 μL pyridine. This solution was mixed by shaking at room temperature for 5 hours. To remove unreacted carboxymethoxylamine, the solution was dried, leaving behind a salt. This was resuspended in 1 mL ethyl acetate and 1 mL DI water, vortexed, and then centrifuged for 1 minute at 2000 x g until the ethyl acetate and water separated from each other. The water phase was discarded, an additional 1 mL of DI water was added, and the process was repeated for a total of three washes. After discarding the final wash, the ethyl acetate solution was dried and the resultant solid was weighed. We recovered 61.6 mg of corticosterone with carboxyl linker. At this stage, the powder contained corticosterone with one to two carboxyl groups conjugated at different locations. The corticosterone-CMO was then conjugated onto the bait-DNA sequence using the EDC/NHS reaction described above. The resulting corticosterone-conjugated DNA was buffer exchanged into DI water using a 3 kDa MWCO Amicon Ultra-0.5 centrifugal filter, and then purified using HPLC and validated for correct bait conjugation via MS.

### DNA-bait synthesis with ATTO 643

Alkyne-bearing DNA oligos (bait-DNA-alkyne4 and bait-DNA-alkyne10) were synthesized using an Expedite 8909 DNA synthesizer at 1 μmol scale. Coupling efficiency during synthesis was monitored after removal of the DMT 5-OH protecting group. After deprotection with a 1:1 aqueous ammonium hydroxide:methyl amine (AMA) solution for 15 minutes at 65 °C, DMT-ON purification of DNA oligos was performed using Glen-Pak DNA purification cartridges following the standard manufacturer’s procedure. Before use, the purified DNA was quantified based on absorbance at 260 nm. The terminal amines of these alkyne-bearing DNA strands were then modified with cortisol or corticosterone following the same protocol as for the Cy5-containing strands. ATTO643 azide was coupled to the cortisol- or corticosterone-modified strands using copper-catalyzed azide-alkyne cycloaddition as described previously(*36*). Briefly, we prepared ∼20 μM DNA and ∼150 μM ATTO 643 azide in 100 mM PBS (pH 7) and added premixed 500 μM BTTES ligand and 100 μM copper (II) sulfate, along with 5 mM sodium ascorbate. This mixture was incubated for 1 hour at room temperature, then purified using a 3 kDa MWCO Amicon Ultra-0.5 centrifugal filter followed by a Glen Gel-Pak 0.2 Desalting Column to exchange the labeled DNA into DI water. Finally, the ATTO 643-labeled DNA was purified using HPLC and validated for correct bait and dye conjugation using MS.

### Synthesis of the antibody-switch

For this first step in the antibody-switch preparation process, ∼250 μg of antibodies were modified as directed in the SiteClick antibody azido modification kit to enzymatically attach azide groups to glycosylation sites in the Fc region of the antibody, yielding a final volume of ∼100 μL 10 μM antibody. Then, 200 mM methyltetrazine DBCO in DMF was added to this ∼100 μL azide-modified antibody to achieve a final concentration of 6.7 mM methyltetrazine DBCO. This step converts the azide into a methyltetrazine group for more efficient click modification with the DNA scaffold. This mixture was incubated for 2 hours at room temperature and then purified with a 50 kDa MWCO Amicon Ultra-0.5 centrifugal filter to remove all free methyltetrazine DBCO. TCO-NHS (axial) was dissolved to 15 mM in anhydrous DMF and then added to scaffold-DNA in 100 mM sodium bicarbonate (pH 9) to achieve final concentrations of 66 μM DNA and 3.75 mM TCO-NHS. This mixture was incubated with rotation overnight at room temperature, then purified with ethanol precipitation and resuspended in DI water to ∼100 μM before use. MS was used to confirm DNA modification. The methyltetrazine-modified antibody and TCO-modified scaffold-DNA were then mixed to achieve a final DNA concentration of 40 μM (and a minimum 3:1 DNA:Ab excess) in 1x PBS. This mixture was incubated overnight at room temperature, then purified with a 50 kDa MWCO Amicon Ultra-0.5 centrifugal filter to remove any free DNA. The degree of labeling was quantified with reducing SDS-PAGE by comparing the intensity of bands from unmodified antibody heavy chains to the intensity of heavier bands formed from specific DNA conjugation to the antibody heavy chains (**Supplementary Figure 2a, b**). The complete removal of free DNA was assessed by denaturing DNA gel electrophoresis, noting the absence of bands corresponding to free scaffold-DNA and the formation of a heavy band corresponding to the antibody-DNA conjugate (**Supplementary Figure 2c**). These gels were prepared by diluting either antibody-DNA or reference scaffold-DNA samples to ∼400 nM in 1x TBE buffer, then mixing in a 1:1 ratio with 95% formamide loading dye(*37*) and heating to 95 °C for 2 minutes. A ladder sample was prepared by diluting low-molecular-weight DNA ladder to 1x in a 1:1 mixture with formamide loading dye, and then heating to 95 °C for 2 minutes. 4 μL of each sample was loaded onto Novex TBE-Urea Gels (10%) and run for 40 minutes at 180 V in 1x TBE buffer before staining with GelStar Nucleic Acid Gel Stain for 5 minutes and imaging.

Splinted ligation was used to covalently attach the bait-DNA conjugate to the antibody-DNA conjugate. First, 3.5 μM bait-DNA was annealed with 3 μM scaffold-complement DNA in ligation buffer (50 mM Tris-HCl (pH 7.5), 10 mM MgCl_2_, and 1 mM ATP) by preparing a 17 μL reaction volume, heating the mixture to 95 °C and then cooling to 4 °C for 30 minutes. Then, 40 μL of ∼500 nM antibody-DNA conjugate was mixed with the entire annealed mixture in ligation buffer and incubated for 1 hour at room temperature to allow for hybridization. 56.3 μL of this mixture was then combined with 13.5 μL of ligation solution, comprising 2,700 U T4 ligase (6.75 μL of 400,000 U/mL) in ligation buffer, and incubated for 1 hour at room temperature to achieve complete ligation into a covalently linked antibody-switch molecule. The success of this ligation was monitored using reducing SDS-PAGE. For the SDS-PAGE analyses mentioned here and above, these products (along with unmodified control antibodies) were diluted to ∼0.2 mg/mL in 8.1 μL of 1x PBS, and then mixed with 0.9 μL of NuPAGE sample-reducing agent and 3 μL of 4X NuPAGE sample loading buffer and heated at 95 °C for 5 minutes. Then, we loaded 10 μL of SeeBlue Pre-stained Protein Standard as a ladder and 12 μL of the antibody mixtures onto NuPAGE 4 to 12% Bis-Tris mini protein gels and ran them for 35 minutes at 200 V with 1x NuPAGE MES SDS running buffer. The gels were stained for 30 minutes with SimplyBlue SafeStain, then destained for up to 2 hours in water before imaging. An increase in the molecular weight of all bands corresponding to DNA-modified heavy chains indicated that all DNA present on the antibody-DNA conjugate had been ligated. This ligation was specific and was not observed in controls where either T4 ligase or the scaffold-complement were not added (**Supplementary Figure 3**).

Finally, the ligated antibody-switches were modified with Alexa Fluor 546 NHS Ester. 15 mM Alexa Fluor 546 NHS ester in DMF was added to 145 nM antibody-switch in 1x PBS to a final fluorophore concentration of 60 μM dye; this reaction was adjusted to pH 9 with 1 M sodium bicarbonate. This mixture was incubated at room temperature for 1 hour, then quenched by adding 33% by volume of 200 mM Tris-HCl (pH 7.5). This resulted in a final antibody-switch concentration of ∼110 nM.

The antibody-switches contain a site-specific biotin moiety within the DNA scaffold and thus can be directly immobilized onto streptavidin-coated surfaces. For each replicate flow cytometry experiment, immobilization was achieved by suspending 1 μL of streptavidin coated Dynabeads T1 in 500 μL of 14.5 nM antibody-switch in physiological buffer. For the fiber-optic biosensor experiments, the streptavidin-functionalized fiber sensor tip was immersed in 40 μL of 75 nM antibody-switch in physiological buffer within the measurement chamber. In both cases, the sensor surfaces were incubated for 1 hour at room temperature, washed with physiological buffer, and then incubated in 5 mM BS3 crosslinker in physiological buffer (500 μL for bead-based assays, 40 μL for fiber-based assays) for 30 minutes to covalently link the antibody-switches to the sensor surface. The beads or fibers were then washed again and incubated in physiological buffer with 5 mM MgCl_2_ and 200 nM scaffold-complement DNA (500 μL for bead-based assays, 40 μL for fiber-based assays) for 2 hours at room temperature, then overnight at 4 °C. Scaffold-complement was added to ensure that the DNA scaffold assumes a double-stranded form. Finally, the beads or fiber sensor were washed and resuspended in physiological buffer (500 μL for bead-based assays, 40 μL for fiber-based assays) before testing.

### Flow cytometry testing of antibody-switch binding, reversibility, and kinetics

Antibody-switch binding responses were measured in physiological buffer or 0.45-μm-filtered chicken plasma. For each replicate curve within a binding curve experiment, a sample of antibody-switch beads (500 uL, prepared as described above) was diluted to 2.8 mL of buffer or plasma per replicate binding curve, then 194 µL of the resulting bead stock was mixed with 6.6 μL of 100 mM digoxigenin or cortisol in DMSO, yielding samples containing 0–3.3 mM target. These samples were incubated for 1 hour at room temperature to equilibrate before analysis on a cell sorter. To monitor FRET signaling, antibody-switches were excited using a 561-nm laser; donor fluorophore (Alexa Fluor 546) emission was measured using one bandpass filter (583 nm center wavelength, 30 nm bandpass width) and acceptor fluorophore (Cy5 or ATTO 643) emission was measured using another bandpass filter (665 nm center wavelength, 30 nm bandpass width). All samples were run to record 3,000-10,000 bead counts.

To measure the reversibility of the switch responses, we tested the response of switch-coated beads over three cycles of target addition and washing. Thus, to study the switch behavior over these cycles, we tested the beads under seven conditions: before adding target, after target addition 1, after wash 1, after target addition 2, after wash 2, after target addition 3, and after wash 3. We recorded three replicates of this series of seven conditions. To prepare each replicate, a sample of antibody-switch-coupled beads (500 µL, prepared as described above) was diluted into 1.5 mL physiological buffer and then split into seven different 200 μL samples to be tested under each of the seven conditions. Switches were subjected to alternating 30-minute cycles of incubation at room temperature in physiological buffer containing either 0 or 1 mM digoxigenin/cortisol. To control for the bleaching or degradation of the antibody-switch-coated bead surface over many cycles of removing and adding new wash or target solution to the beads, we ensured that all seven conditions underwent the same number of solution removal and addition steps, while varying the number and sequence of wash and target steps to achieve the seven desired conditions. All samples were subjected to a single removal and addition of solution at each time step, with the identity of the solution added (either target-free buffer or 1 mM target) specified in **Supplementary Table 2**.

To measure the on-rate kinetics of switching using flow cytometry, a sample of antibody-switch-coated beads (500 µL, prepared as described above) was diluted to 3 mL in physiological buffer and then split into three different 1 mL replicates which were then swapped into 1 mL of buffer containing 1 mM target to measure their kinetic response. To measure the off-rate response, an identical set of bead samples was prepared but then equilibrated in 1 mL of 1 mM target in buffer, followed by swapping them into target-free buffer and subsequently measuring their kinetic response. In both cases, the switch response was measured at 2-to-5-minute intervals after changing concentrations with no additional washes.

### Processing and normalization of binding data

All flow cytometry data were processed in FlowJo 10. Bead populations corresponding to singlet beads were selected, and the median fluorescence of each channel within this population was recorded. All data analysis was performed in Microsoft Excel and GraphPad Prism 8.0.2. All flow cytometry data were corrected for background signal by subtracting fluorescence measurements obtained from unmodified beads run on the flow cytometer at the same settings. For all FRET-based flow cytometry data, donor fluorescent emission (*F*_Donor,_) and acceptor fluorescent emission (*F*_*Acceptor*_) vs. target concentration ([Target]; corresponds to either DIG or Cortisol depending on dataset), the FRET ratio for each datapoint was calculated as:

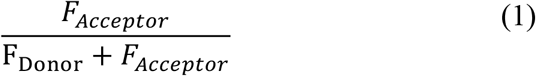

Flow cytometry data were normalized to correct for experiment-to-experiment variations in fluorescent intensity due to changes in instrument gain, bead surface density, and photobleaching between replicates as well as to allow easier visual comparison between different conditions and switches. Normalization was only done by setting a single point to a value of 1, rather than forcing the entire binding curve to span a range of 0 to 1, to maintain a quantitative comparison of the dynamic range and signal of each experiment. Non-normalized FRET ratios and raw FRET data (donor and FRET channels independently) are provided in **Supplementary Figures 4 and 5**. Normalization was carried out per-replicate for all binding curves. For unlinked binding data (**Figures 2b, c** and **4b, c; Supplementary Figure 1)**, binding curves were normalized by setting the largest value in the dataset equal to 1. For antibody-switch binding data (**Figures 2e, 3, and 4d-f**), normalization was done by setting the first FRET measurement in each dataset (either [Target] = 0 or time = 0) to be equal to 1.

### Fitting binding data to obtain affinities, LOD, ULOQ, and kinetic rates

For Cy5-fluorescence based measurements of bait-DNA binding and target competition, we analyzed Cy5 emission (*F*_*Cy*5_) vs. bait-DNA concentration and free target concentration ([Ligand]). These data were fitted to the Langmuir isotherm as follows:

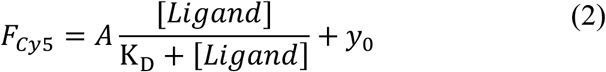

Where K_D_ is the measured affinity of the interaction, and A and *y*_0_ correspond to the magnitude and ligand-free background of the binding response, respectively.

When assessing the antibody-switch binding response, a Hill fit was used to fit the binding data and extract an EC_50_ for each switch, which corresponds to the concentration of target at which 50% of the maximum switching signal is achieved. This fitting was achieved by using non-linear least-squares fitting to the Hill equation:

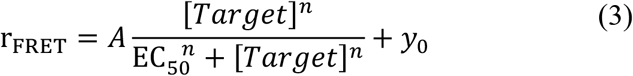

Where n is the Hill coefficient corresponding to the degree of cooperativity observed during switching. K_D_ and EC_50_ values are reported as the best-fit value with 95% confidence interval upper and lower bounds from fitting.

LODs were determined from the mean values of *A, y*_0_, *n*, and EC_50_ as determined from fitting, as well as the standard deviations (*σ*) of these parameters. We applied the standard definition of LOD, which is the target concentration at which the switch response is three standard deviations above background. Thus, LODs for all switches were determined by solving the equation:

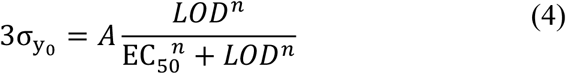

ULOQs were determined from the mean values of *A, y*_0_, *n*, and EC_50_ as determined from fitting, as well as the standard deviations (*σ*) of these parameters. The ULOQ is the concentration at which the switch response is three standard deviations below the saturating signal, which we calculated as follows:

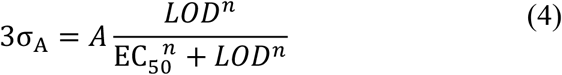

To determine the kinetics of the on- and off-switching response, we first normalized the FRET ratio (r_FRET_) vs. time (t) based on the initial measurement, then fitted these data with a single-phase exponential decay model of the form:

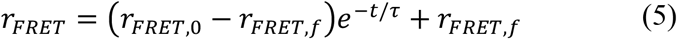

Where *r*_*FRET*,0_ and *r*_*FRET,f*_ are respectively the FRET ratios before and after prolonged laser excitation, and *τ* is the time constant that characterizes the kinetics of antibody-switching behavior.

### Fiber optic measurement system setup

The fiber optic measurement system comprises an optical readout system and tapered fiber optic probe. The development of both components has been thoroughly documented and characterized for continuous sensing in previous work(*31*). Briefly, the optical readout system is designed to excite donor fluorophores on the probe and collect the resulting fluorescent donor and acceptor emission signals, enabling continuous FRET ratio measurements. Excitation light is provided by a 532 nm laser that passes through laser-line filters into a fiber switch. The fiber switch connects either to the sample probe or a dead-end port, allowing measurements with an interval controlled by an Arduino and LabVIEW GUI. When connected, laser light excites the sample probe, then emission light is coupled back into the reader where it is split into two channels using filters to isolate either donor or acceptor emission. These signals are measured using a single photon counting module (SPCM) with an integration time of 100 ms and ∼300 nW output laser power.

### Preparation of fiber optic tip probes

Fiber optic probes were fabricated as described previously(*31*). Briefly, one end of a graded index multi-mode fiber (Thorlabs GIF625) was heated and pulled with a Vytran GPX3000 optical fiber processor (Thorlabs) to form tapered fiber probes with a ∼10-μm tip and an exposed glass surface. The other end of the fiber is terminated by a standard connector that interfaces with the optical readout system. The exposed glass tip was functionalized using a home-built micro-stage setup to dip controlled lengths of the fiber tip into the various solutions needed for functionalization. We cleaned and hydroxyl-activated the fiber optic tips with piranha solution (25% H_2_O_2_ and 75% concentrated H_2_SO_4_) for 2 hours, followed by 2-3 washes with ultrapure water and acetone. The fiber tips were incubated in Vectabond (3% v/v in acetone) for 10 min to functionalize the surface with amine groups, then washed with ultrapure water and dried for 5 min at room temperature. We then coated this surface with a PEG monolayer containing ∼2% biotin-terminated PEG for robust attachment of antibody-switches. This was done by incubating the amine-functionalized surface with a mixture of 32% w/w mPEG-SVA (MW 5,000) and 0.5% w/w biotin-PEG-SVA (MW 5,000) in 0.11 M sodium bicarbonate for 3 hours and then rinsing with water. PEG-functionalized fibers were then placed in 50 μL flow chambers (Grace Biolabs HybriWell Hybridization Cover-HBW6L) mounted onto glass slides to form small measurement chambers with two ports for adding samples during continuous experiments. These chamber-mounted fibers were then stored dry at 4 °C for up to a week before use. Before measurement, the fiber surfaces were incubated with 40 μL of 0.2 mg/mL (∼200 nM) streptavidin solution for 10 min, then washed three times with physiological buffer. The CSO-DNA switch was then immobilized onto the fiber sensor surface as described above. Finally, the switch-functionalized fiber optic probe was connected to the optical readout system for continuous measurements.

### Fiber optic measurement and data analysis

Continuous fiber-based measurements of the antibody-switch response were performed by measuring the switch response for 5 seconds (50 points with 100 ms integration time each) at 30-second intervals. During continuous measurement, the sample concentration within the chamber was varied by pipetting samples with differing cortisol concentration into the chamber at noted time intervals. Samples were injected as two 50 μL injections at each time point to ensure full replacement of the fluid within the chamber. After changing the sample within the chamber, the fiber sensor was left in the new sample with no agitation or flow while recording the donor and acceptor emission and using equation (1) to calculate the corresponding FRET ratio. Intermittently (every ∼1 hour) the laser output power was tested and adjusted to maintain ∼300 nW. To correct for the impact of photobleaching, we first performed a ∼15-minute photobleaching calibration experiment to extrapolate the shift in baseline due to photobleaching over our entire measurement time (**Supplementary Figure 7**). We then subtracted the measured FRET ratio from this predicted FRET baseline to obtain the final “inverted” (signal-ON as opposed to signal-OFF) plots shown in **Figure 5**.

The kinetics of the antibody-switch response were investigated by first isolating the response of the antibody-switch sensor to all individual sample injections during measurement. Each sensor response from the time of each injection to the time of the next injection was separated, then normalized in signal from 0 to 1, where 0 was the smallest signal value and 1 was the largest. These data are shown in **Supplementary Figure 8a** and **b**. These normalized data were then fit with a single exponential as described above for extracting the rate constants of equilibration as a function of final cortisol concentration during equilibration. These rate constants are plotted in **Supplementary Figure 8c**. The lack of dependence of the rate constant on the final concentration is consistent with our proposed antibody-switch mechanism, where both on- and off-switching are limited by the kinetics of slow dissociation events (**Supplementary Discussion 1**).

## Supporting information

Supplementary Figures, Tables, and Discussion

## Acknowledgements

This work was supported by the Chan-Zuckerberg Biohub, the Helmsley Trust and the Wellcome LEAP SAVE program. I.A.P.T. was supported by the Medtronic Foundation Stanford Graduate Fellowship and the Natural Sciences and Engineering Research Council of Canada (NSERC, 416353855). A.A.H. acknowledges support from the Sanjiv Sam Gambhir—Philips Fellowship Program in Precision Health and the NSERC Postdoctoral Fellowships (PDF, Canada). A.P.C. acknowledges support from the NSF Graduate Research Fellowship Program and the Stanford Graduate Fellowship. We thank Dr. Yujia Sun for her assistance with oligonucleotide synthesis and Dr. Tuan Trinh for his assistance with HPLC purification of DNA. This work was supported by the Vincent Coates Foundation Mass Spectrometry Laboratory, Stanford University Mass Spectrometry (RRID:SCR_017801) utilizing the Bruker Microflex MALDI TOF mass spectrometer (RRID:SCR_018696).

## Author contributions

I.A.P.T., J.P., and H.T.S. devised the initial concept. I.A.P.T. devised the design and synthesis of the digoxigenin-responsive antibody-switch. I.A.P.T, J.S., and L.Z. designed and synthesized the cortisol-responsive antibody-switch. I.A.P.T and J.S. designed, executed, and analyzed data from flow cytometry experiments for the digoxigenin and cortisol switches. I.A.P.T., A.H., N.M., and A.C. designed, executed, and analyzed data from fiber optic real-time sensing experiments. I.A.P.T. and H.T.S. wrote the manuscript. J.S., A.H., J.P., and M.E. edited and discussed the manuscript. All authors approved the manuscript.

## References

1. S. S. Gambhir, T. J. Ge, O. Vermesh, R. Spitler, Toward achieving precision health. Sci. Transl. Med. 10, 3612 (2018).

2. T. L. Gruenewald, T. E. Seeman, C. D. Ryff, A. S. Karlamangla, B. H. Singer, Combinations of biomarkers predictive of later life mortality. Proc. Natl. Acad. Sci. U. S. A. 103, 14158–63 (2006).

3. D. H. Hellhammer, S. Wüst, B. M. Kudielka, Salivary cortisol as a biomarker in stress research. Psychoneuroendocrinology. 34, 163–171 (2009).

4. M. Vettoretti, G. Cappon, G. Acciaroli, A. Facchinetti, G. Sparacino, Continuous Glucose Monitoring: Current Use in Diabetes Management and Possible Future Applications. J. Diabetes Sci. Technol. 12, 1064–1071 (2018).

5. J. Kim, A. S. Campbell, B. E.-F. de Ávila, J. Wang, Wearable biosensors for healthcare monitoring. Nat. Biotechnol. 37, 389–406 (2019).

6. A. Kumar, D. Roberts, K. E. Wood, B. Light, J. E. Parrillo, S. Sharma, R. Suppes, D. Feinstein, S. Zanotti, L. Taiberg, D. Gurka, A. Kumar, M. Cheang, Duration of hypotension before initiation of effective antimicrobial therapy is the critical determinant of survival in human septic shock*. Crit. Care Med. 34, 1589–1596 (2006).

7. T. Keller, T. Zeller, D. Peetz, S. Tzikas, A. Roth, E. Czyz, C. Bickel, S. Baldus, A. Warnholtz, M. Fröhlich, C. R. Sinning, M. S. Eleftheriadis, P. S. Wild, R. B. Schnabel, E. Lubos, N. Jachmann, S. Genth-Zotz, F. Post, V. Nicaud, L. Tiret, K. J. Lackner, T. F. Münzel, S. Blankenberg, Sensitive troponin I assay in early diagnosis of acute myocardial infarction. N. Engl. J. Med. 361, 868–877 (2009).

8. H. C. Ates, J. A. Roberts, J. Lipman, A. E. G. Cass, G. A. Urban, C. Dincer, On-Site Therapeutic Drug Monitoring. Trends Biotechnol. (2020), doi:10.1016/j.tibtech.2020.03.001.

9. I. B. Hirsch, E. E. Wright Jr., Using Flash Continuous Glucose Monitoring in Primary Practice. Clin. Diabetes. 37, 150–161 (2019).

10. W. H. Polonsky, D. Hessler, K. J. Ruedy, R. W. Beck, for the DIAMOND Study Group, The Impact of Continuous Glucose Monitoring on Markers of Quality of Life in Adults With Type 1 Diabetes: Further Findings From the DIAMOND Randomized Clinical Trial. Diabetes Care. 40, 736–741 (2017).

11. I. a. Walker, M. Newton, A. t. Bosenberg, Improving surgical safety globally: pulse oximetry and the WHO Guidelines for Safe Surgery. Pediatr. Anesth. 21, 825–828 (2011).

12. C. Chen, Q. Xie, D. Yang, H. Xiao, Y. Fu, Y. Tan, S. Yao, Recent advances in electrochemical glucose biosensors: a review. RSC Adv. 3, 4473–4491 (2013).

13. A. Jubran, Pulse oximetry. Crit. Care. 19, 272 (2015).

14. K. W. Plaxco, H. T. Soh, Switch-based biosensors: A new approach towards real-time, in vivo molecular detection. Trends Biotechnol. 29, 1–5 (2011).

15. Y. Xiao, A. A. Lubin, A. J. Heeger, K. W. Plaxco, Label-Free Electronic Detection of Thrombin in Blood Serum by Using an Aptamer-Based Sensor. Angew. Chem. Int. Ed. 44, 5456–5459 (2005).

16. J. S. Swensen, Y. Xiao, B. S. Ferguson, A. A. Lubin, R. Y. Lai, A. J. Heeger, K. W. Plaxco, H. Tom. Soh, Continuous, Real-Time Monitoring of Cocaine in Undiluted Blood Serum via a Microfluidic, Electrochemical Aptamer-Based Sensor. J. Am. Chem. Soc. 131, 4262–4266 (2009).

17. B. S. Ferguson, D. A. Hoggarth, D. Maliniak, K. Ploense, R. J. White, N. Woodward, K. Hsieh, A. J. Bonham, M. Eisenstein, T. E. Kippin, K. W. Plaxco, H. T. Soh, Real-time, aptamer-based tracking of circulating therapeutic agents in living animals. Sci. Transl. Med. 5 (2013), doi:10.1126/scitranslmed.3007095.

18. N. Nakatsuka, K.-A. Yang, J. M. Abendroth, K. M. Cheung, X. Xu, H. Yang, C. Zhao, B. Zhu, Y. S. Rim, Y. Yang, P. S. Weiss, M. N. Stojanović, A. M. Andrews, Aptamer–field-effect transistors overcome Debye length limitations for small-molecule sensing. Science. 362, 319–324 (2018).

19. P. Dauphin-Ducharme, K. Yang, N. Arroyo-Currás, K. L. Ploense, Y. Zhang, J. Gerson, M. Kurnik, T. E. Kippin, M. N. Stojanovic, K. W. Plaxco, Electrochemical Aptamer-Based Sensors for Improved Therapeutic Drug Monitoring and High-Precision, Feedback-Controlled Drug Delivery. ACS Sens. 4, 2832–2837 (2019).

20. B. Wang, C. Zhao, Z. Wang, K. A. Yang, X. Cheng, W. Liu, W. Yu, S. Lin, Y. Zhao, K. M. Cheung, H. Lin, H. Hojaiji, P. S. Weiss, M. N. Stojanović, A. J. Tomiyama, A. M. Andrews, S. Emaminejad, Wearable aptamer-field-effect transistor sensing system for noninvasive cortisol monitoring. Sci. Adv. 8, 967 (2022).

21. D. Wu, C. K. L. Gordon, J. H. Shin, M. Eisenstein, H. T. Soh, Directed Evolution of Aptamer Discovery Technologies. Acc. Chem. Res. 55, 685–695 (2022).

22. Z. Tang, P. Mallikaratchy, R. Yang, Y. Kim, Z. Zhu, H. Wang, W. Tan, Aptamer Switch Probe Based on Intramolecular Displacement. J. Am. Chem. Soc. 130, 11268–11269 (2008).

23. F. Ricci, A. Vallée-Bélisle, A. J. Simon, A. Porchetta, K. W. Plaxco, Using Nature’s “Tricks” To Rationally Tune the Binding Properties of Biomolecular Receptors. Acc. Chem. Res. 49, 1884–1892 (2016).

24. N. Maganzini, I. Thompson, B. Wilson, H. T. Soh, Pre-equilibrium biosensors as an approach towards rapid and continuous molecular measurements. Nat. Commun. 13, 7072 (2022).

25. M. Kadmiel, J. A. Cidlowski, Glucocorticoid receptor signaling in health and disease. Trends Pharmacol. Sci. 34, 518–530 (2013).

26. L. Thau, J. Gandhi, S. Sharma, “Physiology, Cortisol” in StatPearls (StatPearls Publishing, Treasure Island (FL), 2022; http://www.ncbi.nlm.nih.gov/books/NBK538239/).

27. R. C. Bhake, V. Kluckner, H. Stassen, G. M. Russell, J. Leendertz, K. Stevens, A. C. E. Linthorst, S. L. Lightman, Continuous Free Cortisol Profiles—Circadian Rhythms in Healthy Men. J. Clin. Endocrinol. Metab. 104, 5935–5947 (2019).

28. R. V. Carsia, “Chapter 26 - Adrenals” in Sturkie’s Avian Physiology (Sixth Edition), C. G. Scanes, Ed. (Academic Press, San Diego, 2015; https://www.sciencedirect.com/science/article/pii/B9780124071605000269), pp. 577–611.

29. J. G. Lewis, C. J. Bagley, P. A. Elder, A. W. Bachmann, D. J. Torpy, Plasma free cortisol fraction reflects levels of functioning corticosteroid-binding globulin. Clin. Chim. Acta. 359, 189–194 (2005).

30. G. L. Hammond, Plasma steroid-binding proteins: primary gatekeepers of steroid hormone action. J. Endocrinol. 230, R13–R25 (2016).

31. A. A. Hariri, A. P. Cartwright, C. Dory, Y. Gidi, S. Yee, K. X. Fu, K. Yang, D. Wu, I. A. P. Thompson, N. Maganzini, T. Feagin, B. E. Young, B. H. Afshar, M. Eisenstein, M. Digonnet, J. Vuckovic, H. T. Soh, Continuous optical detection of small-molecule analytes in complex biomatrices (2023), p. 2023.03.03.531030,, doi:10.1101/2023.03.03.531030.

32. J. Das, S. Gomis, J. B. Chen, H. Yousefi, S. Ahmed, A. Mahmud, W. Zhou, E. H. Sargent, S. O. Kelley, Reagentless biomolecular analysis using a molecular pendulum. Nat. Chem. 13, 428–434 (2021).

33. D. Kang, S. Sun, M. Kurnik, D. Morales, F. W. Dahlquist, K. W. Plaxco, New Architecture for Reagentless, Protein-Based Electrochemical Biosensors. J. Am. Chem. Soc. 139, 12113–12116 (2017).

34. C. H. Hansen, D. Yang, M. A. Koussa, W. P. Wong, W. P. W. Designed, Nanoswitch-linked immunosorbent assay (NLISA) for fast, sensitive, and specific protein detection, doi:10.1073/pnas.1708148114.

35. R. Dorey, C. Piérard, S. Shinkaruk, C. Tronche, F. Chauveau, M. Baudonnat, D. Béracochéa, Membrane Mineralocorticoid but not Glucocorticoid Receptors of the Dorsal Hippocampus Mediate the Rapid Effects of Corticosterone on Memory Retrieval. Neuropsychopharmacology. 36, 2639–2649 (2011).

36. S. I. Presolski, V. P. Hong, M. G. Finn, Copper-Catalyzed Azide–Alkyne Click Chemistry for Bioconjugation. Curr. Protoc. Chem. Biol. 3, 153–162 (2011).

37. Cold Spring Harb. Protoc., in press, doi:10.1101/pdb.rec073510.

