## Supplementary Figures, Tables, and Discussion for "An Antibody-Based Molecular Switch for Continuous Biosensing"

### Supplementary Information: An Antibody-based Molecular Switch for Continuous Real-Time Biosensing

**Supplementary Discussion 1 – Thermodynamic and kinetic models of the three-state antibody-switch equilibrium**

The binding properties of the antibody-switch can be modeled as a three-state conformational selection equilibrium that is commonly used to describe the function of completive molecular switches in biosensing(*1*, *2*) (**Figure 1c**). In the absence of target, there is an equilibrium between the closed and open states of the switch. The closed fraction depends on the affinity of the bait molecule for the antibody, $K_D^{bait} = \frac{k_{off}^{bait}}{k_{on}^{bait}}$, where $k_{on}^{bait}$ and $k_{off}^{bait}$ are the on- and off-rates of antibody-bait binding, as well as $C_{eff}$, which is the effective concentration of the bait experienced by the antibody as a result of their covalent tethering. Importantly, $C_{eff}$ does not represent a concentration, but rather the probability of binding interactions occurring between the linked antibody-bait pair. In the absence of free target, the ratio of closed:open switches is $\frac{[C_{eff}]}{K_D^{bait}}$:1. Introduction of free target molecules leads to antibody binding with $K_D^{target} = \frac{k_{off}^{target}}{k_{on}^{target}}$, where $k_{on}^{target}$ and $k_{off}^{target}$ are the on- and off-rates of antibody-target binding, shifting the equilibrium away from the closed state and towards the bound state. This switching leads to fluorescent signaling, yielding an overall effective sensor dissociation constant of:

$$K_D^{eff} = K_D^{target}\left(1 + \frac{[C_{eff}]}{K_D^{bait}}\right) \tag{4}$$

This effective dissociation constant governs the sensitivity of the switch, and this relationship demonstrates how substitution of the bait molecule can manipulate the overall affinity of the switch. By keeping switch assembly the same (*i.e.,* unchanged $C_{eff}$) while varying the bait identity to weaken its antibody binding (higher $K_D^{bait}$), we can decrease $K_D^{eff}$ overall, yielding a higher-affinity switch. We note that $K_D^{bait}$is equivalent to the concentration at which the antibody-switch signal reaches half maximal value, ($EC_{50}$), which we have used later when applying a Hill fit to our data; this trend between increasing $K_D^{bait}$ and decreasing $K_D^{bait}$/$EC_{50}$ holds in that case.

To understand the kinetics of antibody-switch sensing, we can break down each of the transitions of the three-state equilibrium into their constituent forward and reverse binding reaction rates (**Supplementary Figure 9**). The bait or target on-rates ($k_{on}^{bait}\left[C_{eff}\right]$ or $k_{on}^{target}[T]$) in this equilibrium will be fast relative to the off-rates at reasonable free or effective target concentrations. For example, assuming concentrations of 10 nM to 100 µM target and an antibody on-rate consistent with literature-reported rates(*3*) of $10^5$ $M^{-1}s^{-1}$, we find on-rates of $10^{-3}$ to $10^1$ $s^{-1}$. This is similar to or higher than representative antibody off-rates, which range from $10^{-3}$ to $10^{-5}$ $s^{-1}$. Thus, the process of switching, which occurs through transitions between the closed and bound states, will be limited by the slow dissociation reactions with rate constants $k_{off}^{target}$ or $k_{off}^{bait}$. Thus, because the bait is selected to have partial identity to the target molecule, the overall observed switching rates when exposed to concentration increases or decreases will be similar, as we have observed in our measurements.

#### Supplementary Figures

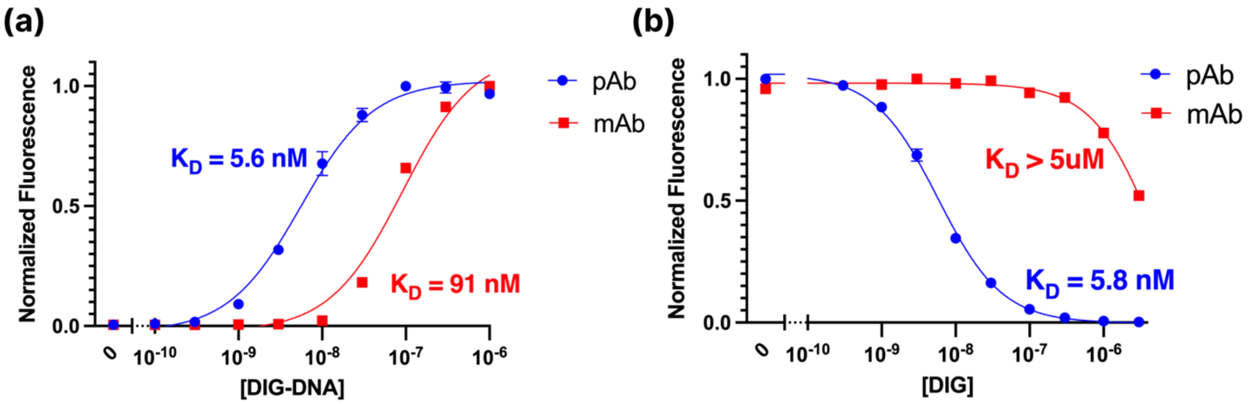


**Supplementary Figure 1 | Antibody binding to DIG-DNA. (a)** Flow cytometry-based measurement of DIG-DNA binding affinity to either a monoclonal (mAb, red) or polyclonal (pAb, blue) candidate antibody shows substantially weaker affinity for the former. **(b)** We assessed competitive binding between DIG-DNA and free DIG to both antibody candidates with varying free DIG concentrations and either 10 nM DIG-DNA for pAb or 50 nM DIG-DNA for mAb, to compensate for weaker affinity. These results show that the monoclonal exhibits poor molecular competition and thus is poorly-suited for this antibody-switch.

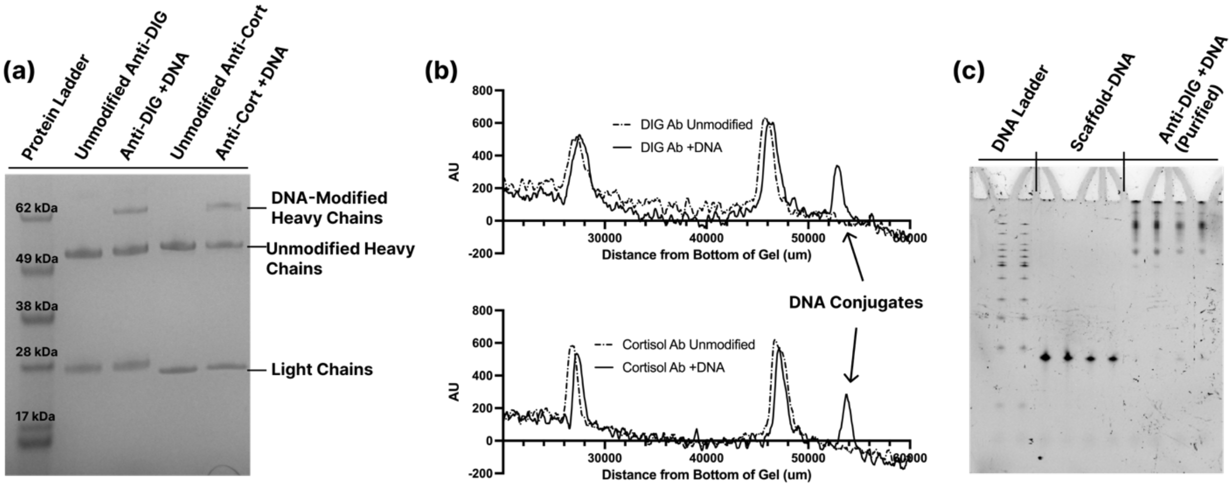


**Supplementary Figure 2 | Validation of site-specific antibody-DNA conjugation. (a)** Reducing SDS-PAGE comparing unmodified and DNA-conjugated antibodies for both digoxigenin (Anti-DIG) and cortisol (Anti-Cort) shows selective DNA conjugation to the antibodies' heavy chains. **(b)** Quantitative analysis of the SDS-PAGE for both targets shows that a substantial proportion of the heavy chains are DNA-conjugated. While the yield is not 100%, only conjugated antibodies contain the biotin needed for surface immobilization, such that unconjugated antibodies do not interfere with sensor preparation. **(c)** Denaturing DNA gel electrophoresis indicates successful purification of our antibody-DNA conjugates and removal of excess unconjugated DNA.

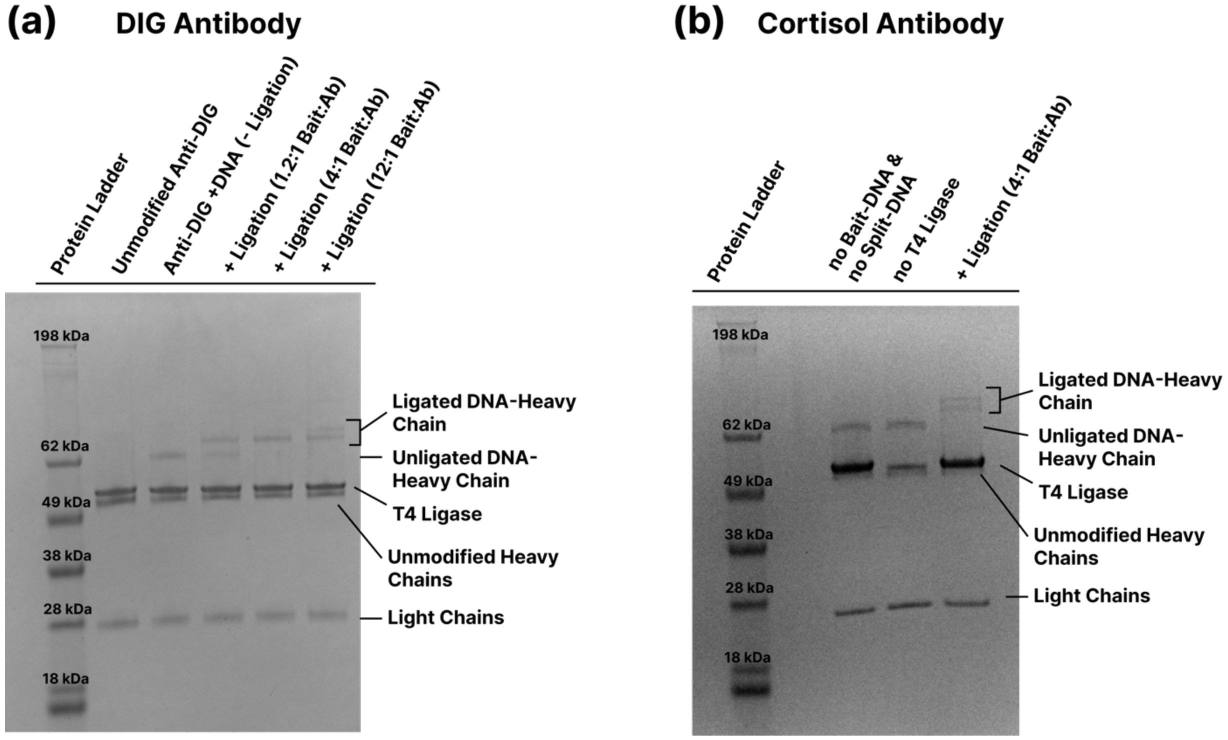


**Supplementary Figure 3 | Validation of the ligation step for switch assembly. (a)** SDS-PAGE analysis of ligation with T4 ligase for the anti-digoxigenin antibody and DIG-DNA bait with a variety of bait:antibody ratios. Successful ligation leads to an increase in molecular weight for the DNA-conjugated heavy chain bands, while all other bands remain unmodified. This ensures that only biotinylated antibodies (via the scaffold-DNA) are further modified to incorporate the bait. At a ratio of 4:1, we observe complete modification, and this ratio was used for all subsequent syntheses. **(b)** The same ligation process for the anti-cortisol antibody-switch. Only the full ligation reaction with 4:1 bait:antibody results in the disappearance of unligated heavy chain-DNA conjugates and the formation of heavier, ligated products.

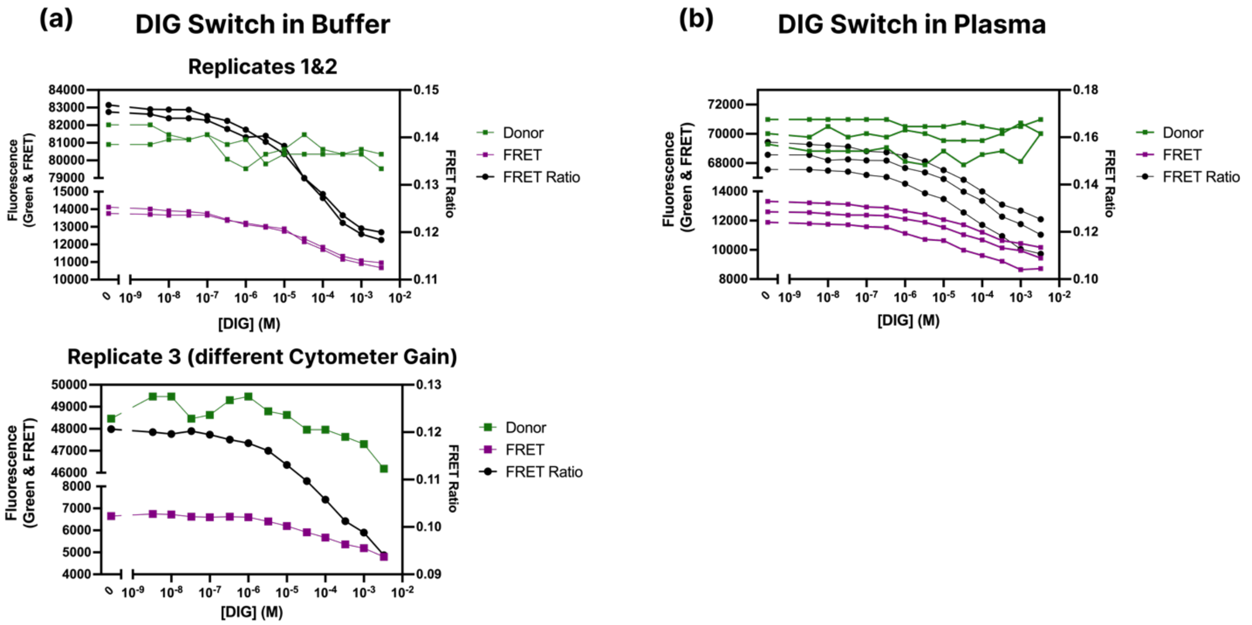


**Supplementary Figure 4 | Raw FRET Data for the DIG-responsive antibody-switch. (a)** Raw measurements of donor emission (green) and FRET emission (purple) as well as the corresponding un-normalized FRET ratio (black) for three replicate experiments using the DIG antibody-switch in buffer. The third replicate (bottom) was performed at different gain settings, which leads to a scaling of the values but not of the overall affinity or sensitivity of the switch. Once normalized for gain differences, the replicates align well, as seen in **Figure 2e. (b)** Raw FRET measurements from flow cytometry for the DIG antibody-switch in undiluted chicken plasma, with data shown for three replicate experiments.

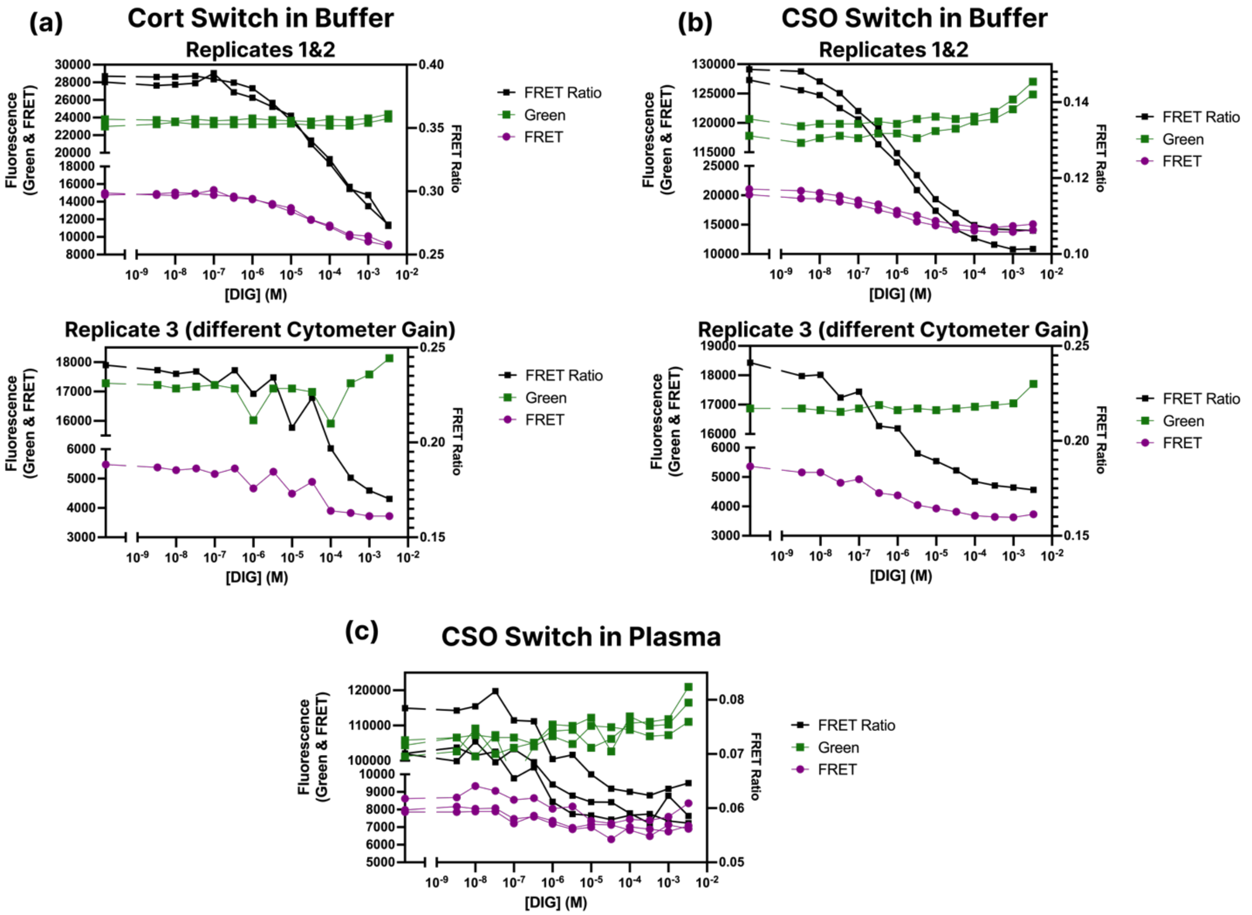


**Supplementary Figure 5 | Raw FRET data for cortisol-responsive switches. (a)** Raw measurements of donor emission (green) and FRET emission (purple) as well as the corresponding non-normalized FRET ratios (black) for the Cort-DNA antibody-switch in buffer. **(b)** Raw FRET measurements from flow cytometry for the CSO-DNA antibody-switch in buffer. For both **a** and **b**, the bottom panels show a third replicate experiment performed with different cytometer gain, as described in **Supplementary Figure 4. (c)** Raw FRET measurements from flow cytometry for the CSO-DNA antibody-switch in undiluted chicken plasma.

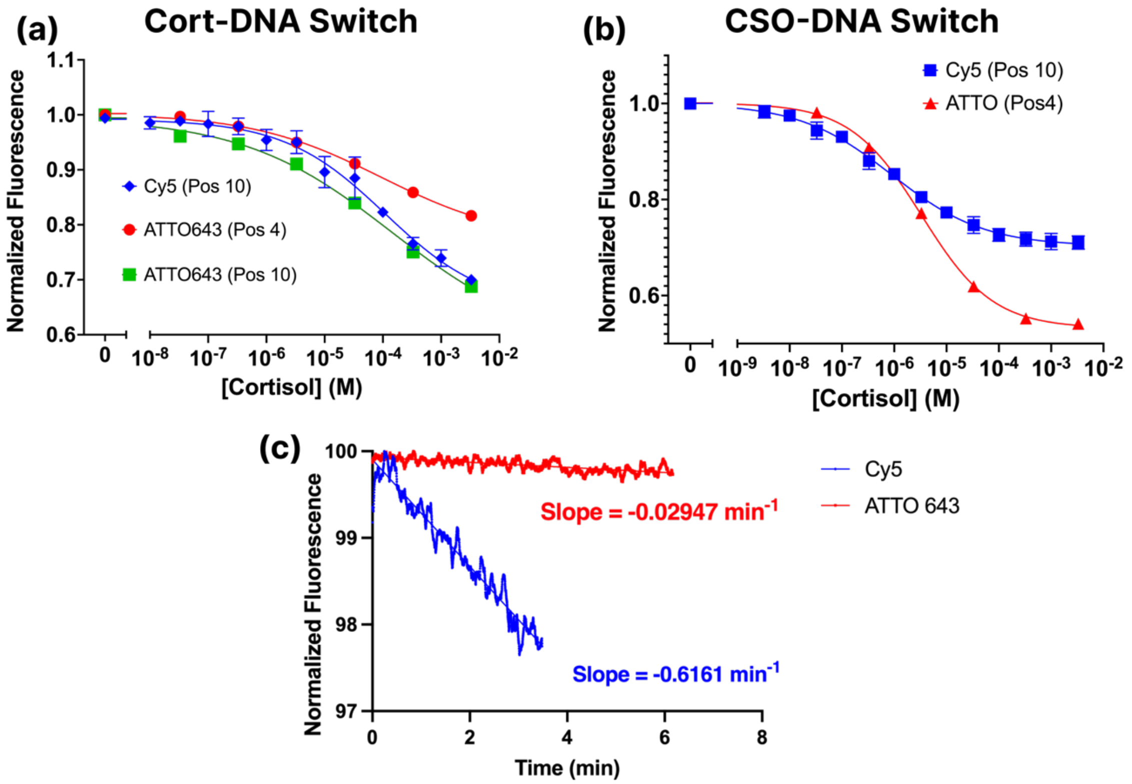


**Supplementary Figure 6 | ATTO643-based switch performance relative to Cy5. (a)** Flow cytometry testing of the cortisol response of antibody-switches constructed using the Cort-DNA bait with different acceptor fluorophore identities or locations. The switch responds to cortisol similarly regardless of whether the acceptor is Cy5 or ATTO 643, or of its position within the DNA sequence (Pos4 is 4 bases from the bait attachment, Pos10 is 10 bases away). **(b)** Flow cytometry testing of cortisol-responsive antibody-switches constructed with CSO-DNA indicates that acceptor dye idenitity and position have a minimal impact on the switch function. **(c)** Continuous measurement of the switch in target-free buffer on a fiber-optic sensor at ~300 nW excitation laser power shows that the ATTO 643 design is much more resistant to photobleaching compared to the Cy5-based design.

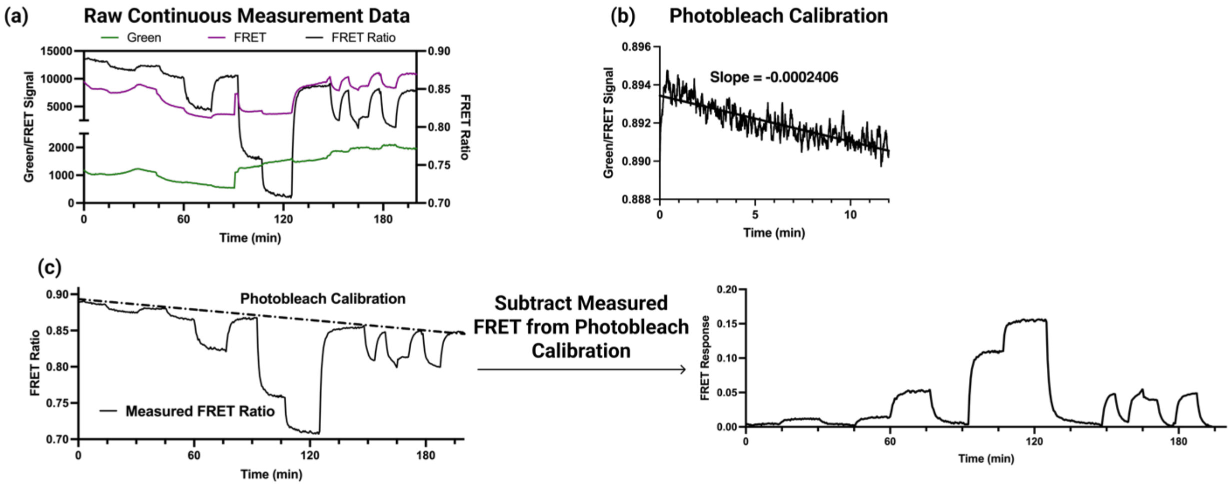


**Supplementary Figure 7 | Photobleaching correction and processing of fiber probe data. (a)** Raw measurements of donor emission (green) and FRET emission (purple) as well as the corresponding un-normalized FRET ratio (black) from the fiber optic antibody-switch sensor over the course of a >3 hour measurement. Discontinuities in the green and FRET channels result from adjustment of the laser to maintain constant excitation power in cases of laser drift, but these discontinuities do not affect the FRET ratio measurements. **(b)** We corrected for photobleaching over the duration of continuous measurement by first measuring the sensor FRET decay due to acceptor photobleaching in the absence of target. **(c)** While photobleaching leads to a slow decay in FRET ratio over the course of the >3-hour experiment, our calibration data predict this drop, thus allowing us to cancel the photobleaching-induced signal changes by subtracting the measured ratio from the extrapolated photobleaching decay rate. This also produces an inversion of the signal (FRET ratio drops appear as increases), which allows for more intuitive visualization of the real-time measurement data.

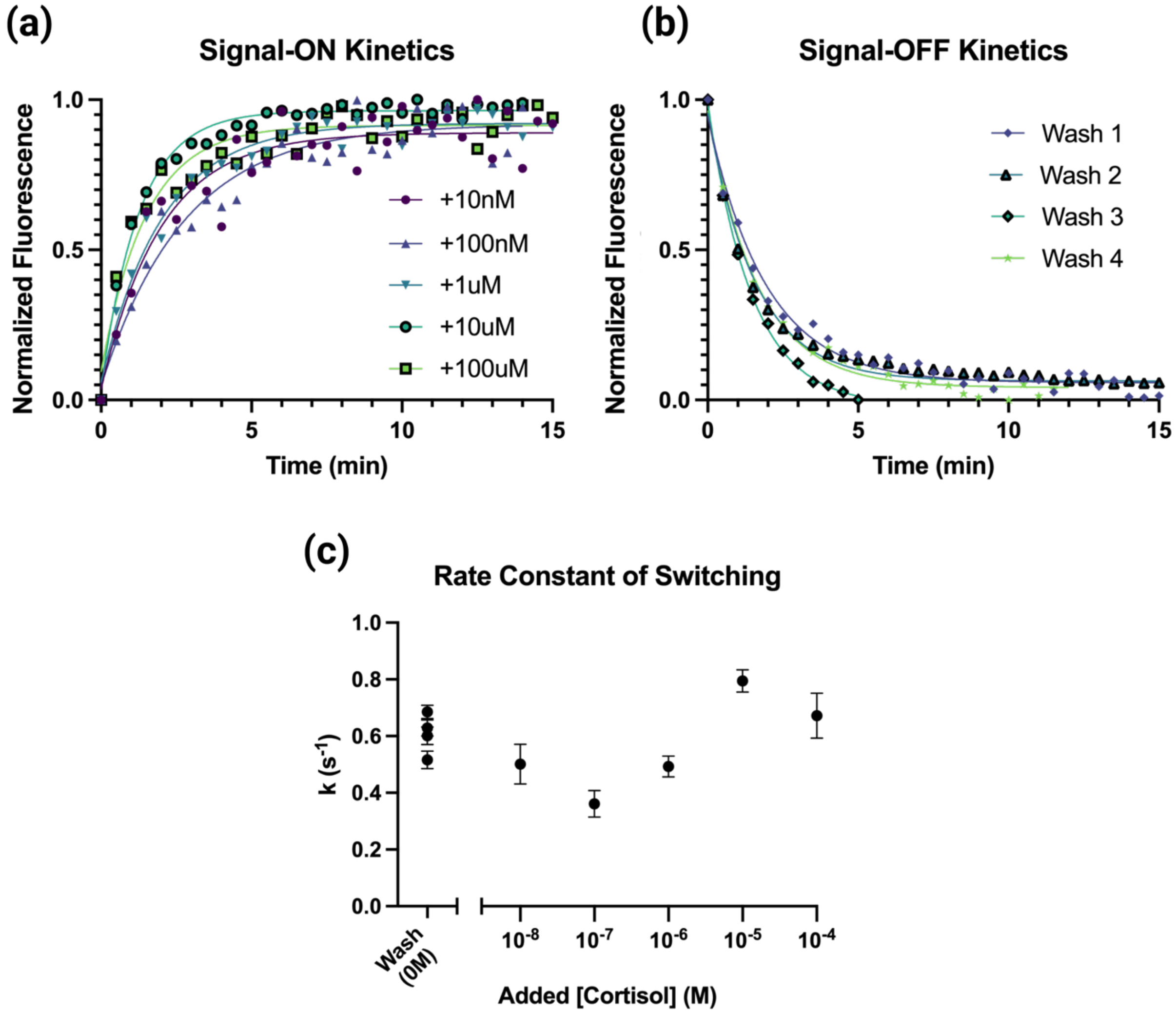


**Supplementary Figure 8 | Kinetics of continuous antibody-switch sensing. (a)** Normalized sensor responses to the injection of increasing cortisol concentrations into the measurement chamber over three hours of continuous measurement, where labels indicate the final injected concentration. At each concentration, the sensor equilibrates within five minutes to changing concentrations. **(b)** Normalized sensor responses to the injection of cortisol-free buffer into the measurement chamber during our three-hour continuous measurement. With each wash, sensor equilibration occurs within 5 minutes. **(c)** Comparing the antibody-switch response rate-constants for both decreasing (wash) or increasing final concentrations shows that the rate constant is independent of the concentration. These rate constants correspond to sensor equilibration time-constants in the range $\tau = 1.3$ to $2.8\ min^{-1}$. Thus, the switch achieves < 5 minute equilibration regardless of the concentration.

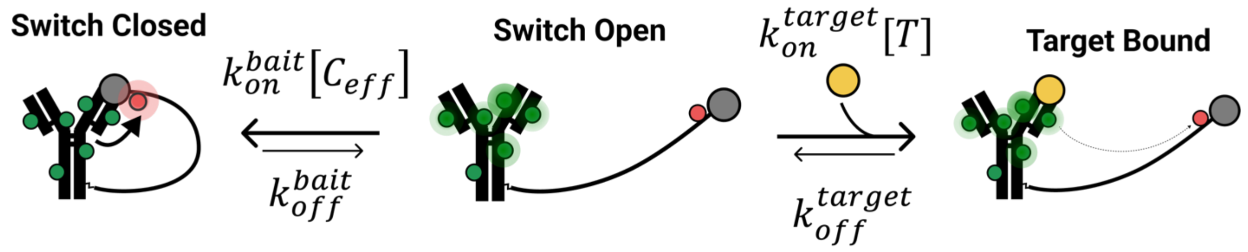


**Supplementary Figure 9 | Kinetic model of antibody-switch binding.** Transitions between the three states of the antibody-switch binding model depend on the on- and off-rates of antibody binding to the bait or target molecule, as well as the concentration of free target and the effective concentration of bait ($C_{eff}$) experienced by the antibody due to tethering of the bait. Importantly, the relatively slow off-rates limit transitions from closed to bound and vice versa, leading to roughly symmetric switching kinetics.

**Supplementary Table 1 – DNA sequences used in this work**

| Sequence Name | Sequence (5'-3') |
|---|---|
| Bait-DNA | /5AmMC6/GGCAGAGGCA/iCy5/TTTTCGAGC |
| Bait-DNA-Alkyne4 | /1*/GG CAG AGG CA/2*/ TTT TCG AGC |
| Bait-DNA-Alkyne10 | /1*/GG C/2*/G AGG CAT TTT TCG AGC |
| Scaffold-DNA | /5Phos/CAGTAATAAGAGAATATAAAGTACCGACAAAAGG /iBiodT/AAAGTA/3AmMO/ |
| Scaffold-Complement (ligation splint) | TACTTTACCTTTTGTCGGTACTTTATATTCTCTTATTACTG GCTCGAAAAATGCCTCTGC |
| **Modifications** | |
| /5AmMC6/ | 5' amino modifier C6 from IDT |
| /iCy5/ | Int Cy5 from IDT |
| /5Phos/ | 5' phosphorylation from IDT |
| /iBiodT/ | Int biotin dT from IDT |
| /3AmMO/ | 3' amino modifier from IDT |
| /1*/ | 5'-amino-modifier C6 from Glen Research |
| /2*/ | Alkyne-modifier serinol phosphoramidite from Glen Research |

**Supplementary Table 2 – Sequence of target addition and wash cycles used for experiments to assess reversibility of bead-coupled antibody-switch activity.**

Each column describes the series of reagent additions used to test a sample condition in our bead-based reversibility experiment. Cycles corresponding to +Buffer were done by removing the supernatant from the beads and adding target-free buffer, while cycles of +1 mM correspond to removal of the supernatant and addition of 1 mM DIG or cortisol. Each bead condition was subjected to the same number of cycles to control for the effects of supernatant removal and bleaching.

| | **Tested Bead Condition** | | | | | | |
|---|---|---|---|---|---|---|---|
| **Cycle of Reagent Addition** | **Buffer** | **+Target** | **+Wash** | **+Target 2** | **+Wash 2** | **+Target 3** | **+Wash 3** |
| **1** | + Buffer | + Buffer | + Buffer | + Buffer | + Buffer | + Buffer | +1 mM |
| **2** | + Buffer | + Buffer | + Buffer | + Buffer | + Buffer | +1 mM | + Buffer |
| **3** | + Buffer | + Buffer | + Buffer | + Buffer | +1 mM | + Buffer | +1 mM |
| **4** | + Buffer | + Buffer | + Buffer | +1 mM | + Buffer | +1 mM | + Buffer |
| **5** | + Buffer | + Buffer | +1 mM | + Buffer | +1 mM | + Buffer | +1 mM |
| **6** | + Buffer | +1 mM | + Buffer | +1 mM | + Buffer | +1 mM | + Buffer |
| | Measure via flow cytometry | | | | | | |

**Supplementary References**


1. F. Ricci, A. Vallée-Bélisle, A. J. Simon, A. Porchetta, K. W. Plaxco, Using Nature's "Tricks" To Rationally Tune the Binding Properties of Biomolecular Receptors. *Acc. Chem. Res.* **49**, 1884–1892 (2016).

2. B. D. Wilson, A. A. Hariri, I. A. P. Thompson, M. Eisenstein, H. T. Soh, Independent control of the thermodynamic and kinetic properties of aptamer switches. *Nat. Commun.* **10**, 5079 (2019).

3. J. P. Landry, Y. Ke, G.-L. Yu, X. D. Zhu, Measuring Affinity Constants of 1,450 Monoclonal Antibodies to Peptide Targets with a Microarray-based Label-Free Assay Platform. *J. Immunol. Methods*. **417**, 86–96 (2015).
